# Similarity and strength of glomerular odor representations define neural metric of sniff-invariant discrimination time

**DOI:** 10.1101/356279

**Authors:** Anindya S. Bhattacharjee, Sasank Konakamchi, Dmitrij Turaev, Roberto Vincis, Daniel Nunes, Atharva A. Dingankar, Hartwig Spors, Alan Carleton, Thomas Kuner, Nixon M. Abraham

**Author notes:** These authors contributed equally to the work (Anindya S. Bhattacharjee, Sasank Konakamchi, Nixon M. Abraham). Senior and corresponding authors. Deceased on 13th May 2018 (Sasank Konakamchi).

## Abstract

The olfactory environment is first represented by glomerular activity patterns in the olfactory bulb. It remained unclear, how these activity patterns intersect with sampling behavior to account for the time required to discriminate odors. Using different classes of volatile stimuli, we investigated glomerular activity patterns and sniffing behavior during olfactory decision-making. Mice discriminated monomolecular odorants and binary mixtures on a fast time scale and learned to increase their breathing frequency at a fixed latency after trial initiation, independent of odor identity. Relative to the increase in breathing frequency, monomolecular odorants were discriminated within 10-40 ms while binary mixtures required an additional 60-70 ms. Intrinsic imaging of odor-evoked glomerular activity maps in anesthetized and awake mice revealed that the Euclidean distance between glomerular patterns elicited by different odors, a measure of similarity and activation strength, was anti-correlated with discrimination time. Therefore, the similarity of glomerular patterns and their activation strengths, rather than sampling behavior, define the extent of neuronal processing required for odor discrimination, establishing a neural metric to predict olfactory discrimination time.

## Introduction

Sensory inputs are constantly processed by the brain and serve as a basis for making decisions, ultimately driving behavior. The accuracy and speed of decision-making varies depending on the urgency and difficulty in accomplishing a task (Abraham et al., 2004; Reddi and Carpenter, 2000; Rinberg et al., 2006), although this idea has recently been challenged (Zariwala et al., 2013). Currently, the speed of olfactory discrimination is subject of a fruitful debate focusing on a causal link between stimulus-dependency of decision-making (Abraham et al., 2004, 2010, 2012; Kay et al., 2006; Rinberg et al., 2006; Schaefer and Margrie, 2007) and the uncoupling of decision-making accuracy from sampling duration (Uchida et al., 2006; Uchida and Mainen, 2003; Zariwala et al., 2013). In support of the latter proposition, using enantiomers, it has been shown that odor categorization is achieved in a definite temporal window independent of task complexity and urgency (Zariwala et al., 2013). Do these opposing conclusions arise from differences in odorants used? Since previous studies relied on using limited number of odorants, the high dimensionality of olfactory stimulus space was not considered as a possible parameter impacting on decision-making behavior.

Sensory neurons of the olfactory epithelium project in a receptor-specific manner to olfactory bulb (OB) glomeruli (Ressler et al., 1994). Odorants belonging to different chemical classes evoke complex combinatorial patterns of activity among glomeruli (Friedrich and Korsching, 1998; Kauer and White, 2001; Mori et al., 1999). The contribution of spatial coding towards olfactory discrimination becomes crucial taken the fact that an individual glomerular module can be explained as a molecular feature detecting unit (Mori et al., 1999) and has been supported by *in vivo* functional imaging studies from the OB (Abraham et al., 2014; Bathellier et al., 2007; Kauer, 1988; Rubin and Katz, 1999; Spors and Grinvald, 2002; Spors et al., 2006; Uchida et al., 2000; Vincis et al., 2012; Wachowiak and Cohen, 2001; Xu et al., 2003). Monomolecular odorants often evoke glomerular patterns showing little overlap, while their binary mixtures typically activate strongly overlapping glomerular patterns (Abraham et al., 2004; Fletcher, 2011). Mice accurately discriminate monomolecular odorants within hundreds of milliseconds, while discrimination of binary mixtures requires additional tens of milliseconds (Abraham et al., 2004, 2012; Rinberg et al., 2006). These two observations prompted the question, if the similarity of glomerular odor representations is related to observed differences in odor discrimination time (ODT) for different odorants. Such a relationship would establish a simple metric linking similarity of odor representations and extent of neuronal processing required for highly accurate odor discrimination.

Beyond glomerular activity patterns, odor sampling behavior could impact the ODT in an odor-specific manner. Active sampling behavior is critical in encoding different physical features of stimuli present in fluctuating odor plumes. For example, locusts move their antennae actively to sample the odors that may help in localizing the source (Huston et al., 2015); flicking of antennules may help lobsters in sensing the concentration gradient (Koehl et al., 2001) and rodents increase frequency of breathing, referred to as sniffing, while detecting novel odors, engaging in social interactions or discriminating varying concentrations (Jordan et al., 2018b; Shusterman et al., 2018; Wesson, 2013; Wesson et al., 2008a, b). Under different contexts animals make olfactory-driven decisions in a few milliseconds (Abraham et al., 2004, 2010, 2012, 2014; Resulaj and Rinberg, 2015; Rinberg et al., 2006; Uchida and Mainen, 2003; Wesson et al., 2008a), underlining the importance to understand the contribution of odor sampling to the mechanisms defining odor discrimination time.

Here, we investigated how glomerular activation patterns relate to ODT for +/– carvones, +/– octanols, cineol (CI)/eugenol (EU) and isoamyl acetate (IAA)/ethyl butyrate (EB) in addition to the odor pair amyl acetate (AA) and ethyl butyrate studied so far. The choice of odorants was governed by two factors: (1) dorsal location of the activated glomeruli to facilitate functional imaging of odor representations in anesthetized (Abraham et al., 2004; Johnson et al., 2002; Rubin and Katz, 2001) and awake mice; (2) overlapping glomerular activity patterns - enantiomers were selected as they are stereoisomers that are non-superimposable and mirror images of each other (Ma and Shepherd, 2000; Rubin and Katz, 2001; Uchida and Mainen, 2003). Our correlation studies of functional imaging and behavioral measurements revealed that ODT is inversely correlated with the Euclidean distance between glomerular activation patterns across chemical classes and varying concentrations, suggesting that strength and similarity of the patterns govern ODT. Although, the active sampling measured by breathing frequency during the decision-making time window did not show any dependence on the ODT difference between the monomolecular odorants and binary mixtures, we did observe a significant increase in breathing frequency during the decision-making period.

## Results

### Stimulus-dependent olfactory discrimination time in multiple odor pairs

To determine if odorants of different chemical classes exhibit stimulus-dependent ODTs we first trained a group of mice for AA/EB, CI/EU and C+/C– (set 1, see STAR Methods). Following an initial training of 1200 trials on CI/EU, we tested AA versus EB to establish direct comparisons to our previous data (Abraham et al., 2004). Mice reached asymptotic performance levels for AA vs. EB discrimination within 600 trials (Figure 1a). Learning the mixture discrimination took longer but reached 95% accuracy at the end of the block (range: 200-500 trials; Figure 1a and 1b). ODT was determined from the last three hundred trials of each block (Figure 1c). Mice discriminated AA/EB simple odor pair in 255 ± 16 ms (mean ± SEM, n = 10 mice, data averaged over two blocks of 300 trials), but they took significantly more time to discriminate the binary mixtures (359 ± 16 ms, one way RM ANOVA, F = 31.2, p = 6.4 x 10^-9^). These results suggest that ODT is a stable and reproducible behavioral measurement that reflects the similarity of olfactory stimuli.

**Figure 1.**
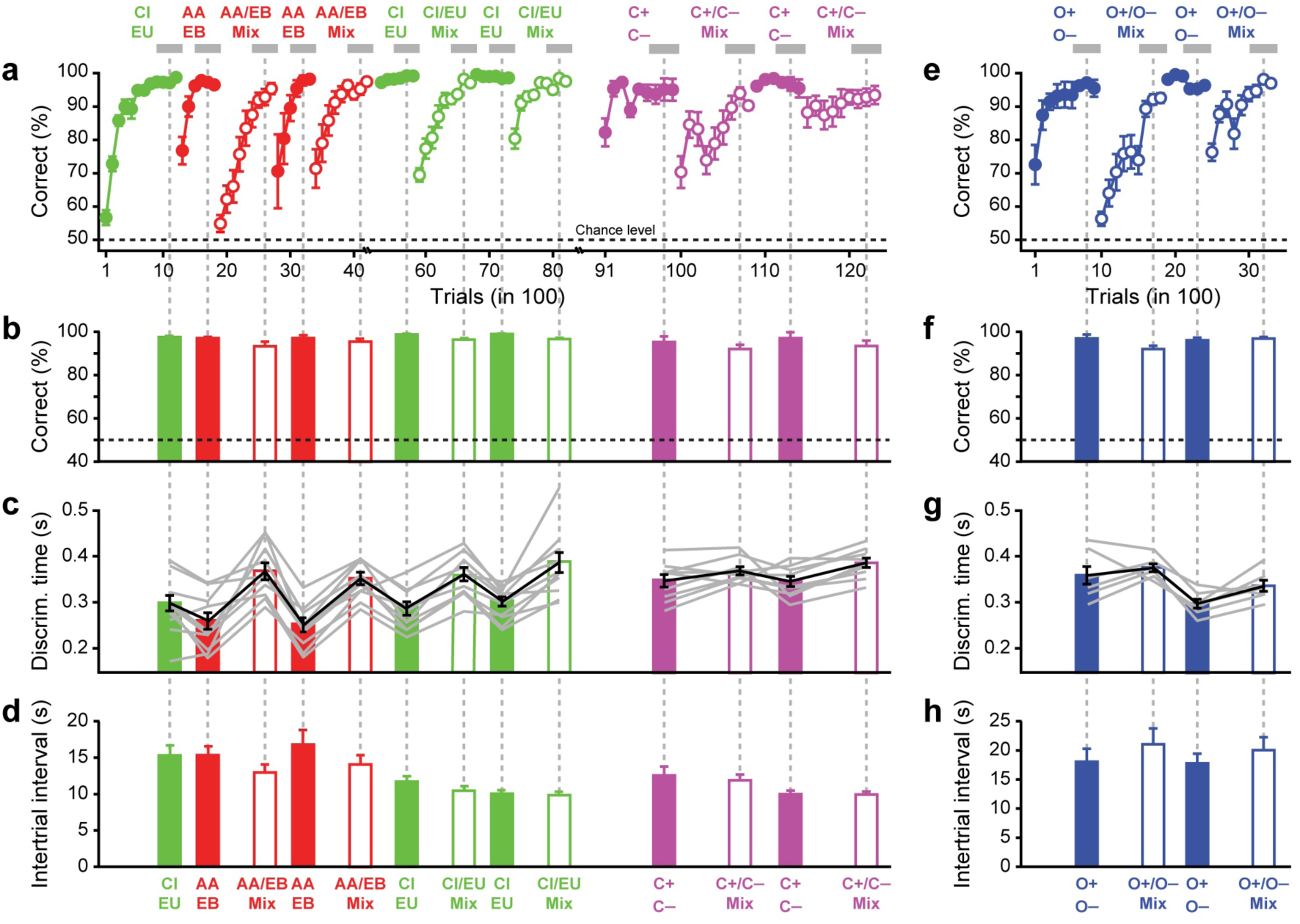
Discrimination time varies for pairs of odors belonging to different chemical classes. (a) Training schedule and accuracies measured for amyl acetate (AA) / ethyl butyrate (EB), cineol (CI) / eugenol (EU), (+)-carvone (C+) / (–)-carvone (C–) and binary mixtures (60-40 vs. 40-60). Accuracy of discrimination shown as % correct choices of 100 trials. Each data point is the average of 9-10 animals (see STAR Methods). The abscissa reflects progression of time. Analysis of ODT restricted to the areas highlighted with a grey bar. Odor pairs used were 1% CI vs. 1% EU, 1% AA vs. 1% EB, 1% C+ vs. 1% C– and the mixtures of these odorants as indicated (60-40 mixtures were used, for example AA-EB mix is 0.6%AA+0.4%EB vs. 0.4%AA+0.6%EB; all odor pairs were counterbalanced as described in STAR Methods section). (b) Accuracy corresponding to experimental blocks indicated in **a** (grey bars). (c) ODTs for individual mice and for the population (grey and, black lines, respectively), averaged for the same period as in **b**, are larger for pairs of binary mixtures than for the pairs of monomolecular odorants. (d) ITI as a measure of motivation is independent of odor similarity. (e) Training schedule for octanols and binary mixtures. Accuracy of discrimination shown as % correct choices of 100 trials. Each data point is the average of 7 animals (different from animals shown in **a**). The abscissa reflects progression of time. Odor pairs used were 0.1% O+ vs. 0.1% O– and mixtures of O+ and O– (0.06% O+ - 0.04% O– vs. 0.04% O+ - 0.06% O–). (f) Accuracy (averaged for the period indicated by grey bars in **e**). (g) ODTs corresponding to experimental blocks indicated in **e**. (h) ITI interval for octanols and binary mixtures tasks. Data are presented as mean ± SEM.

Training was then continued with CI/EU and their binary mixtures. Mice discriminated the monomolecular odorants CI/EU with maximal accuracy (Figure 1a). While the performance dropped transiently upon introduction of binary mixtures, mice still performed well above chance level and performed at >90% success rates during the second episode (Figure 1a and 1b). Discrimination times for mixtures were consistently longer than those determined for the simple odor pair even when the order of discrimination tasks consisting of simple odors and binary mixtures were randomized (Figure 1c, n = 10 mice, one way RM ANOVA, F = 17.5, p = 4.7 x 10^-8^). Training with alternating presentations of simple odors and binary mixtures did not alter performance levels and ODTs at the end of each training session (Figure 1b and 1c). On average, mice discriminated CI/EU simple odor pair in 295 ± 10 ms (mean ± SEM, n = 10 mice; data averaged over three blocks of 300 trials), but they took ∼80 ms more to discriminate the binary mixtures (373 ± 17 ms, data averaged over two blocks of 300 trials).

In this behavioral paradigm, mice have control over the inter-trial interval (ITI, Figure 1d), a parameter reflecting the overall motivation and arousal state of the mice (Abraham et al., 2004, 2010). ITI and ODT for different monomolecular odorants and binary mixtures were, however, not correlated (R^2^ = 0.2, ANOVA F = 1.8, p = 0.2), suggesting that differences in discrimination time are not related to motivational factors or arousal state.

As enantiomers represent a class of challenging stimuli due to their overlapping patterns of activity (Ma and Shepherd, 2000; Rubin and Katz, 2001; Uchida and Mainen, 2003), mice were further trained for C+/C- and their binary mixtures. ODT measured for monomolecular odorants was again significantly faster than for the binary mixtures (345 ± 7 ms and 377 ± 7 ms for C+/C- and mixtures, respectively; mean ± SEM, n = 9 mice, data averaged over two blocks of 300 trials, one way RM ANOVA, F = 3.4, p = 0.03). The ITI did not show any trend to explain the difference in ODT between simple odors and binary mixtures (Figure 1d, R^2^ = 0.1, ANOVA, F = 0.6, p = 0.4).

To consider a possible effect of odor concentration (all odor pairs were tested at 1% dilution in mineral oil and further 20% dilution in air) and to extend the number of odor pairs studied, we tested another enantiomer pair (+)-Octanol/(-)-Octanol (O+/O-) and their binary mixtures at a lower dilution of 0.1%. After pre-training, mice (set 2, see STAR Methods) were first trained for CI/EU, followed by O+/O- and their binary mixtures. Mice acquired simple O+/O- discrimination within 400 trials (Figure 1e). Mixture discrimination required 900 trials to reach a 90% performance level (Figure 1e and 1f), comparable to the previously tested odor pairs (Figures 1a and 1b). While performance dropped transiently upon the second testing of binary mixtures, mice still performed well above chance level and stabilized at > 95% for the last 300 trials of mixture training (Figure 1e and 1f). ODT for the binary mixtures of O+/O- (355 ± 7 ms, mean ± SEM) was significantly increased by ∼30 ms compared to monomolecular odors (327 ± 13 ms, n = 7 mice, data averaged over two blocks of 300 trials; one way RM ANOVA, F = 9.2, p = 6.3 x 10^-4^) (Figure 1g). The ITI did not show any correlation with the difference in ODT measured for simple odors and binary mixtures (Figure 1h, R^2^ = 0.4, ANOVA, F = 1.4, p = 0.4), suggesting that the differences in discrimination time cannot be explained by changes in motivation or activity levels of the animals. These experiments show that mice need more time to discriminate the binary mixtures of O+ and O- than for monomolecular O+ and O- and this increase in ODT measured at a lower odor concentration was comparable to increases in ODTs measured for other odorants at higher concentrations. ODT for enantiomers can be measured reliably across different animals and more time is required for the accurate discrimination of closely related binary mixtures of enantiomers than for monomolecular enantiomers. In summary, our results show that stimulus-dependent ODT is a general property of odor discrimination in mice.

### Extent of stimulus dependency varies for different odor pairs

Monomolecular odorant pairs were discriminated with distinct ODTs across different stimuli classes [Figure 2a, filled bars, in this figure, ODTs are pooled from the freely moving (FM) as well as head-restrained (HR) mice]. While comparing the same number of mice and trials from the very last training session, FM and HR mice did not show significant differences (Figure S1; see STAR Methods). For example, AA/EB could be discriminated much faster than C+/C- and O+/O-. All odor pairs were discriminated with distinct ODTs apart from the enantiomers (Figure 2a, comparison between the monomolecular odorants, ANOVA, F = 11.09, p < 0.0001; Fisher’s LSD: AA/EB vs. CI/EU, p < 0.05, AA/EB vs. C+/C- and O+/O-, p < 0.0001, CI/EU vs. C+/C-, p < 0.01, CI/EU vs. O+/O-, p < 0.05 and C+/C- vs. O+/O-, p = 0.34). Hence, at maximal performance, the ODT is an odorant-specific property with each odorant requiring a distinct time to be discriminated. While an increase in ODT was observed for each of the different binary mixtures tested, the magnitude of the ODT increase differed between the stimuli (Figure 2a, ANOVA, monomolecular odorants vs. binary mixtures: F = 12.81, p < 0.0001, Fisher’s LSD, AA/EB vs. AA-EB 60-40 Mix: p < 0.0001, CI/EU vs. CI-EU 60-40 Mix: p < 0.0001, C+/C- vs. Carvones 60-40 Mix: p < 0.05, O+/O- vs. Octanols 60-40 Mix: p < 0.01). When plotting the increase of ODT of binary mixtures against the ODT of simple odors, the data points could be described with a regression line (Figure 2b). This relation predicts that the increase in ODT between monomolecular odorants and mixtures is inversely proportional to the monomolecular ODT. Furthermore, the intersection of the regression line with the abscissa may define a maximal ODT of approximately 400 ms at maximal discrimination accuracy. If the discrimination of a simple odor pair approaches 400 ms, its binary mixture can be discriminated with the same ODT at maximal accuracy. In conclusion, monomolecular odorants that can be discriminated fast are predicted to require longer ODTs when discriminated as binary mixtures. Accordingly, if monomolecular odorants require near maximal time for discrimination, i.e. ∼400 ms, their binary mixture will be discriminated with no additional consumption of time. This observation may suggest an upper limit of olfactory decision-making time for odor discrimination.

**Figure 2.**
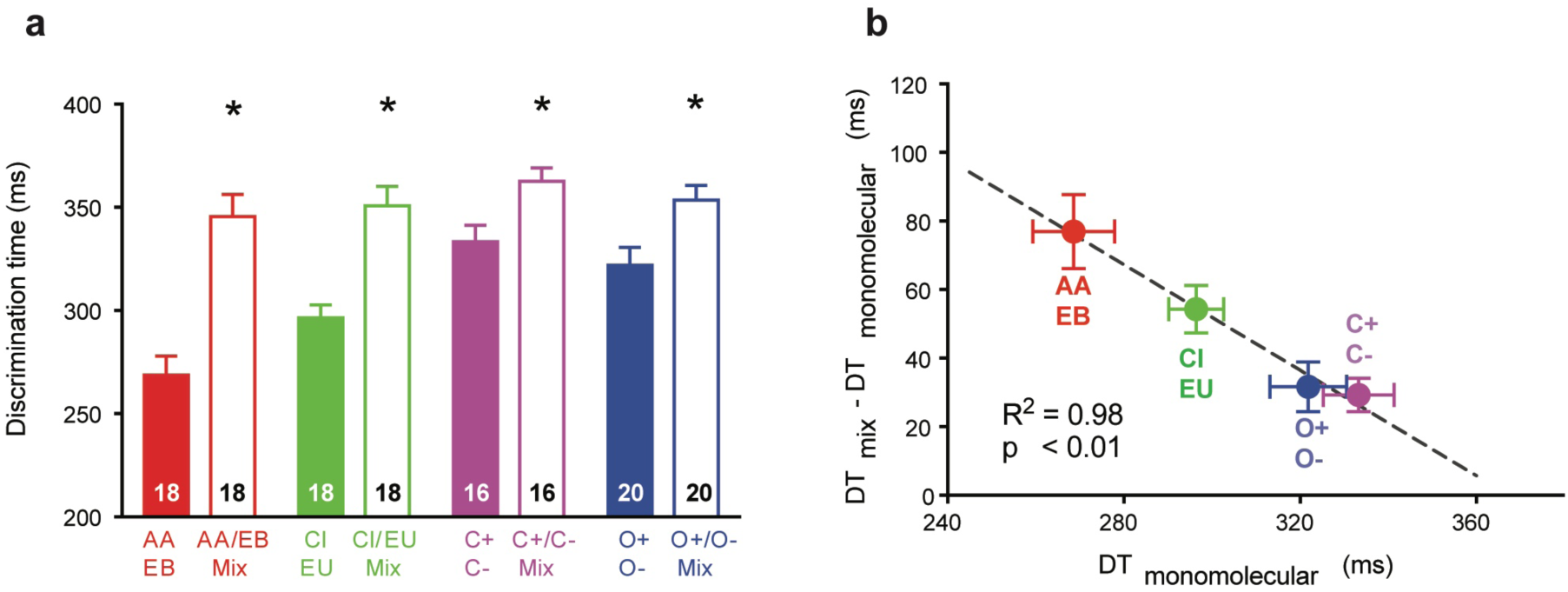
Comparison of ODTs for simple monomolecular odorants and complex binary mixtures across different classes of chemicals. (a) Average ODTs across different experiments. Data are presented as mean ± SEM. DT_AA/EB_ = 269 ± 9 ms, DT_AA/EBmix_ = 346 ± 11 ms, DT_CI/EU_ = 296 ± 6 ms, DT_CI/EUmix_ = 351 ± 9 ms, DT_C+/C_– = 333 ± 8 ms, DT_C+/C–mix_ = 363 ± 7 ms, DT_O+/O–_ = 322 ± 9 ms, DT_O+/O–mix_ = 353 ± 7 ms. Number of animals is indicated on each bar. * Comparison of ODTs for monomolecular odorants versus binary mixtures: Figure 2a, ANOVA, monomolecular odorants versus binary mixtures: F = 12.81, p < 0.0001, * Fisher’s LSD, p < 0.05) (b) Delta ODT for binary mixtures and associated monomolecular odorants is inversely related to ODT for monomolecular odorants (linear regression: R^2^ = 0.98, ANOVA, p < 0.001). Data are presented as mean ± SEM. See also Figure S1

### Sampling behavior does not underlie increase of ODT in mixture discrimination

Our results demonstrated an increase in the ODTs for binary mixtures compared with monomolecular odorants of different chemical classes. Can this difference be caused by the animals’ sampling behavior towards the stimuli of varying complexity? To investigate this question, we trained a batch of naïve mice (set 4, see STAR Methods) on monomolecular odorants and their binary mixtures used in the previous experiments and measured their breathing frequency throughout the olfactory experiment. Breathing was measured non-invasively using a pressure sensor while mice performed odor discrimination tasks under head-restrained conditions (See STAR Methods, Figure S2).

We found ODTs similar to those measured in freely moving mice (Figure S1). The average breathing frequency was 3.9 ± 0.05 Hz during the entire 2 s odor application and 4.5 ± 0.05 Hz during the time window at which the mice performed odor discrimination, here also referred to as the decision-making time period (Figure 3a,b, average ± SEM). Hence, breathing frequency increased significantly during odor sampling, a process typically referred to as sniffing (Figure 3c, comparison between pre-, during and post-decision-making time periods one-way ANOVA, p < 0.05, n = 7-8 mice). Naïve mice trained on a mineral oil (MO) vs. MO task, the solvent used to dilute odorants for the odor discrimination tasks, did not develop episodes of stimulus-related sniffing (one-way ANOVA, p > 0.05, n = 8 mice). Our data also show that breathing frequencies for discrimination of monomolecular odorants and their respective binary mixtures were indistinguishable (Welch’s t-test, p > 0.05), suggesting that they can be excluded as a mechanism contributing to the increased ODT required to discriminate two highly similar stimuli. Lastly, we did not find a difference in breathing frequencies between different chemical classes apart from AA/EB (ANOVA, F = 7.62, p < 0.0001, Fisher’s LSD, comparison of AA/EB and AA-EB 60-40 binary mixture vs. all other chemical classes, both simple and binary mixtures, least p < 0.01). In summary, breathing frequency is modulated independently of odor complexity during the period of odor discrimination and does not contribute to increased ODT.

**Figure 3.**
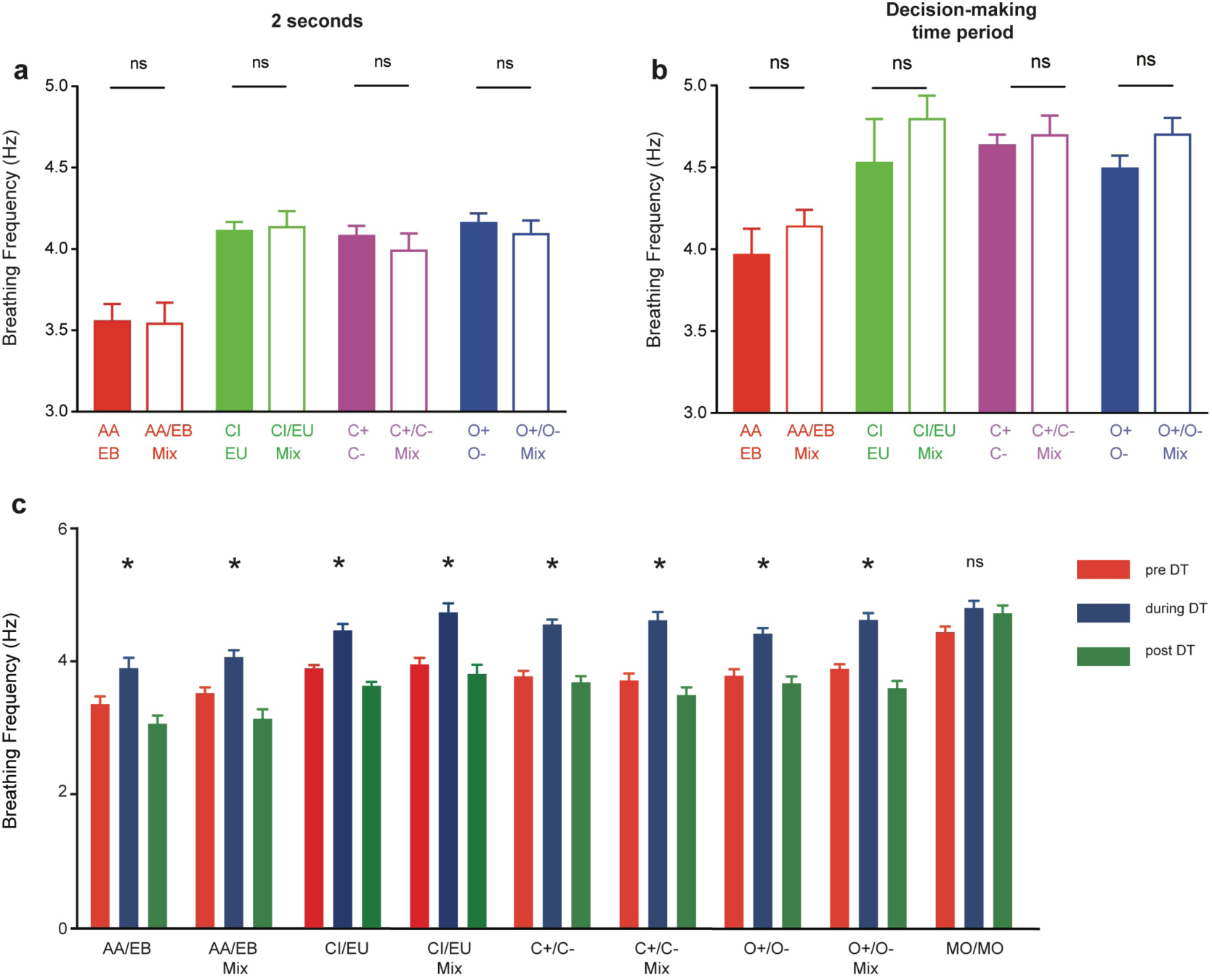
Sniffing behavior during decision-making. (a) Sniffing frequencies (SFs) were measured during the entire stimulus duration i.e. for 2 s. No difference was found in the SFs for each chemical class. Data are presented as mean ± SEM. SF_AA/EB_ = 3.56 ± 0.1 Hz, SF_AA/EBmix_ = 3.54 ± 0.12 Hz, SF_CI/EU_ = 4.11 ± 0.05 Hz, SF_CI/EUmix_ = 4.13 ± 0.09 Hz, SF_C+/C_– = 4.08 ± 0.06 Hz, SF_C+/C–mix_ = 3.99 ± 0.1 Hz, SF_O+/O–_ = 4.15 ± 0.06 Hz, SF_O+/O–mix_ = 4.09 ± 0.08 Hz. Comparison between monomolecular odors and binary mixtures (same odor pairs were used as in Figures 1 & 2, but behavioral training was done under head-restrained conditions) was done using Welch’s t-test, p > 0.05 for all odor pairs tested (n = 7-8 mice). (b) No difference in sniffing frequencies measured during the decision-making time. ODTs and the corresponding SFs were done for each task (300 trials) where the accuracy of performance was ≥ 80%. Data are presented as mean ± SEM. SF_AA/EB_ = 3.97 ± 0.15 Hz, SF_AA/EBmix_ = 4.14 ± 0.1 Hz, SF_CI/EU_ = 4.53 ± 0.1 Hz, SF_CI/EUmix_ = 4.79 ± 0.14 Hz, SF_C+/C_– = 4.63 ± 0.06 Hz, SF_C+/C–mix_ = 4.70 ± 0.12 Hz, SF_O+/O–_ = 4.49 ± 0.08 Hz, SF_O+/O–mix_ = 4.70 ± 0.1 Hz. SFs from the decision-making time periods for monomolecular odors and the corresponding binary mixtures were compared using Welch’s t-test. p > 0.05, for all the odor pairs tested (n = 7-8 mice). (c) SFs calculated during, pre-, and post-decision-making time periods. SF increased during the decision-making period for all odor pairs tested (*Comparison between pre-, during and post-decision-making time periods one-way ANOVA, p < 0.05, n = 7-8 mice). For naïve animals trained on MO/MO discrimination task, we did not observe any significant differences during the three periods (one-way ANOVA, p > 0.05, n = 8 mice). See also Figure S2

### Temporal relationship of sniffing and decision-making

Our data so far suggests that the decision-making time window coincides with an increased breathing frequency, yet sniffing does not seem to account for increases in ODT when analyzing breathing frequencies on a rather coarse scale (Figure 3). To further examine the relationship of sniffing and decision-making, we looked at the temporal profile of changes in breathing frequency during the odor application (Figure 4). The raster plot on the bottom of the panels depicts the start of breathing cycles detected during all trials (see STAR Methods). The raster plot was translated into a histogram using a 20 ms bin width. Evidently, the breathing frequency increased reliably in all experiments at around 250 ms after starting the odor delivery (Figure 4, blue line). To check if this increase is odor-dependent, we trained a naïve batch of mice on MO vs. MO, where the animals expectedly did not learn (Figure S3a, ANOVA, F = 1.66, p = 0.16). Comparison of the MO vs. MO breathing histogram (Figure S3b and grey trace in Figure 4a) with the odor discrimination tasks revealed a highly significant difference (Figure 4, Kolmogorov-Smirnov (K-S) test, p < 0.0001 for all odor pairs), indicating that increased sniffing occurred only when mice were conditioned to discriminate odors. To probe for synchronization of sniffing and licking, we looked at the onset of licks in response to rewarded and non-rewarded odors. The results show that random licking responses preceding the decision-making time window account only for a small average rise, while the peak of lick response occurs after the decision time window. A stereotyped phase relationship between sniffing and licking is not evident (green histogram in Figures 4 & S4, K-S test, p < 0.0001 for all odor pairs).

**Figure 4.**
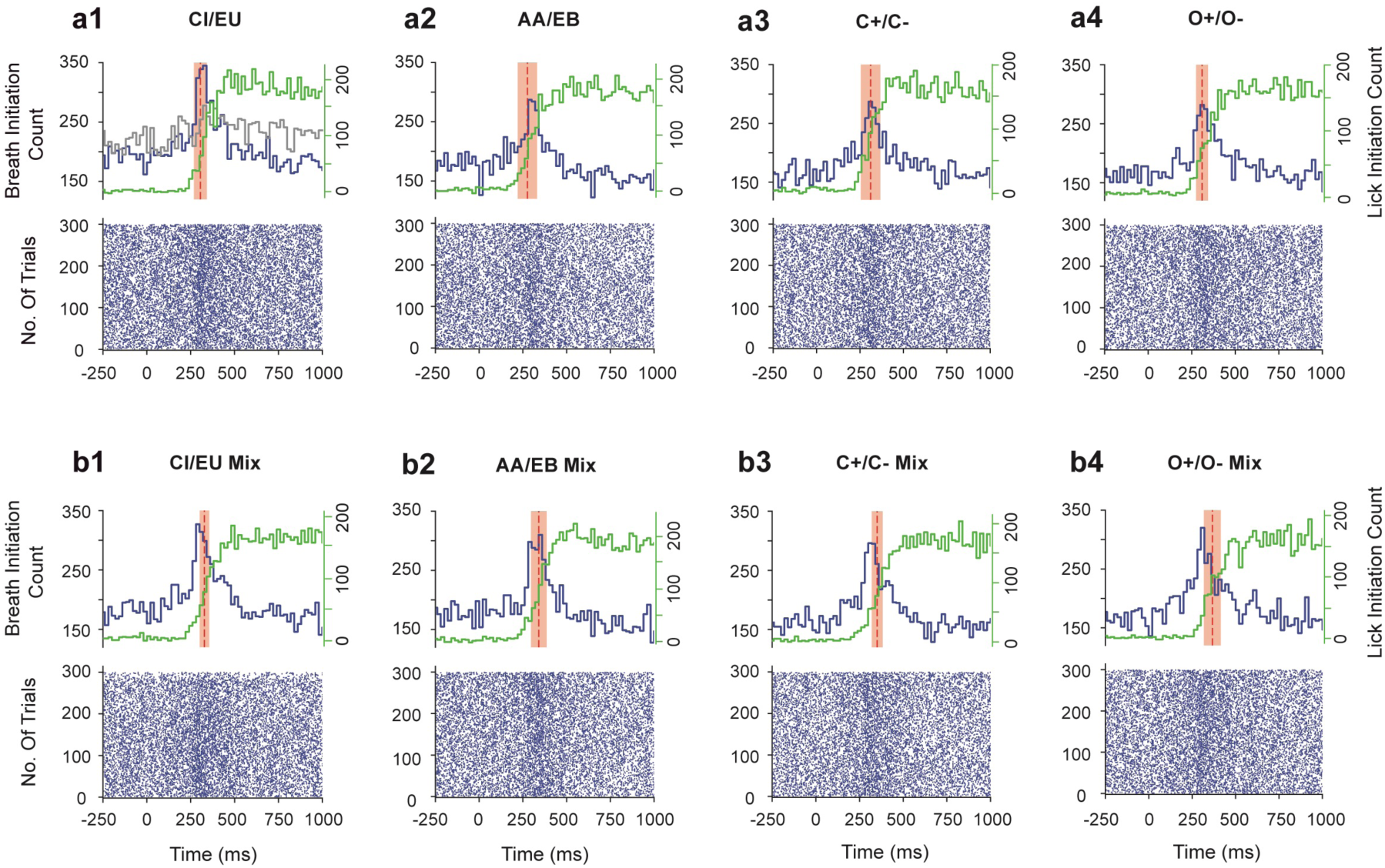
Sniffing, licking, decision-making and ODT under head-restrained conditions. Bottom half of each panel (raster plots of a1 – b4) shows the breathing cycle pattern of the mice during the last task of training where the performance accuracy was ∼90%. Each point on the raster represents the start of inhalation. Top half of each panel (a1 – b4) shows a histogram taken from all trials using a bin size of 20 ms (i.e. adding up the number of breathing cycles initiated within 20 ms across all trials). The red dotted line and shaded region represent the mean ODT ± SEM. The histograms for each odor pair have been compared with the MO vs. MO. The MO histogram trace is shown for comparison in panel a1 (grey line). Comparison between the odor and MO histograms were done using Kolmogorov-Smirnov (K-S) test; p < 0.0001 for all odor pairs. Green traces represent the histogram of licking onset in response to odorants. K-S test comparing Breath Initiation Count histograms and Lick Initiation Count histograms; p < 0.0001 for all odor pairs. See also Figures S3 and S4

Interestingly, Figure 4 reveals that the peak of sniffing coincides with ODT for monomolecular odorant discriminations (stippled red lines indicating mean ODTs and shaded box representing SDs), yet precedes ODT for mixture discriminations. To dissect this further we calculated the ODTs (Figure S5) and the sniffing peak latency (SPL, Figure S6) for each task when mice performed with > 80% accuracy (Out of 122 tasks pooled for simple and complex binary mixture odor pairs, one mouse performed with 76% accuracy for one task of CI/EU and for all other odor pairs and tasks, the performance for the same mouse was >80%). While ODTs increased for binary mixtures, the SPL remained unaltered (Figure 5a, comparison between ODTs for monomolecular odorants and associated binary mixtures: Paired t test, p < 0.05. Comparison between SPL for simple odors and associated binary mixtures: Paired t test, p > 0.05; Figure 5b, cumulative probability, K-S test, p = 0.8, n = 61). The shift for ODTs compared with SPL is evident for binary mixtures (Figure 5d, cumulative probability, K-S test, p < 0.05, n = 61), but not for monomolecular odorants (Figure 5c, cumulative probability, K-S test, p = 0.8, n = 61). The additional time elapsing from sniffing peak latency to ODT amounts to approximately 30 ms (Figure 5e, cumulative probability, K-S test, p < 0.01, n = 61). This time interval may arise from neuronal processing in the olfactory system in preparation to achieve accurate and rapid odor discrimination. An important aspect of our study is that animals could sample the odor according to their individual sniffing strategies. This resulted in a spread of inhalation onsets ranging from the 1^st^ millisecond to almost one respiratory cycle in single trials. Therefore, we calculated the medians task-wise and these vary in the range of 100-200 ms. We calculated the first breath onset latencies for simple and complex odors and we did not observe any differences (Figure S7a). We also calculated the ODTs corrected for these delays and we do see slower ODTs for binary mixtures compared to simple odors, which correspond to additional time intervening from SPL to ODT measurements (Figure S7b).

**Figure 5.**
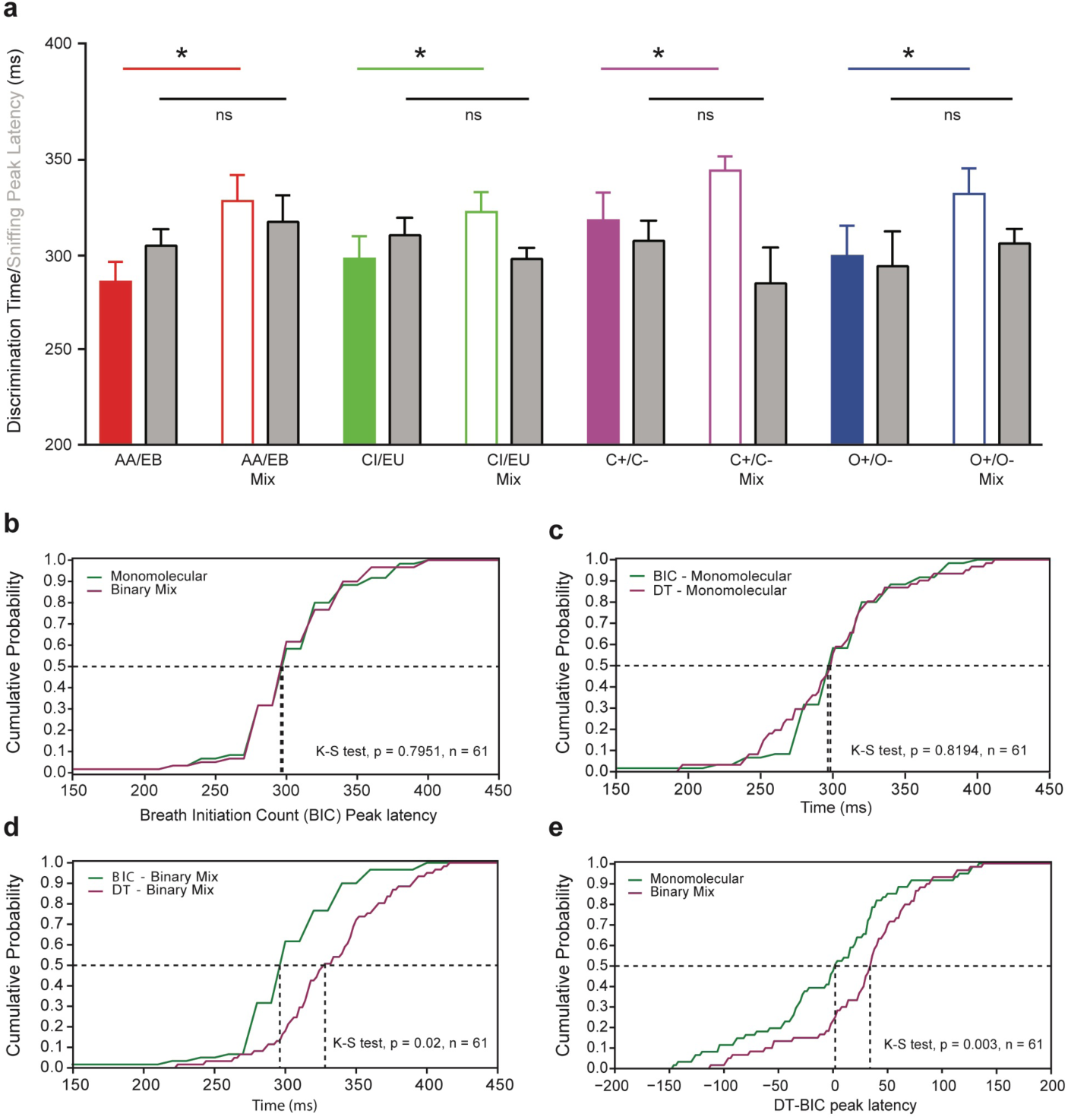
Sniff-invariant odor discriminations for odor pairs of varying complexity. (a) Comparison between ODTs and SPL for odor pairs of varying similarity. Colored (filled and empty) bars are ODTs, grey bars represent the sniffing peak latency values for the corresponding ODTs. Data are presented as mean ± SEM. DT_AA/EB_ = 286 ± 11 ms, DT_AA/EBmix_ = 328 ± 14 ms, DT_CI/EU_ = 298 ± 12 ms, DT_CI/EUmix_ = 322 ± 11 ms, DT_C+/C_– = 318 ± 15 ms, DT_C+/C–mix_ = 344 ± 7 ms, DT_O+/O–_ = 299 ± 16 ms, DT_O+/O–mix_ = 332 ± 13 ms. sniffing P_AA/EB_ = 305 ± 9 ms, sniffing P_AA/EBmix_ = 317 ± 14 ms, sniffing P_CI/EU_ = 310 ± 9 ms, sniffing P_CI/EUmix_ = 298 ± 6 ms, sniffing P_C+/C_– = 307 ± 11 ms, sniffing P_C+/C–mix_ = 285 ± 19 ms, sniffing P_O+/O–_ = 294 ± 18 ms, sniffing P_O+/O–mix_ = 306 ± 8 ms. Comparison between ODTs for monomolecular odorants and associated binary mixtures: Paired t test, p < 0.05. Comparison between sniffing peak latency for simple odors and associated binary mixtures: Paired t test, p > 0.05. (b) The SPL compared between monomolecular odors and binary mixtures by pooling all the odor pairs. The SPL remains unaltered for monomolecular odors and binary mixtures (K-S test, p = 0.8, n = 61) (c) and (d) Comparison between cumulative probability distributions of ODT and SPL for monomolecular odors and binary mixtures. Additional time is required to discriminate binary mixtures (compared with SPL, 5d, K-S test, p = 0.02, n = 61), but, ODTs and SPL remain unaltered for monomolecular odors (5c, K-S test, p = 0.82, n = 61). (e) The difference between SPLs and ODTs for monomolecular odors and binary mixtures represented by cumulative probability distributions. The difference is greater for binary mixtures as compared to monomolecular odors (K-S test, p = 0.003, n = 61). See also Figures S5, S6 and S7

### ODT is correlated to strength and similarity of odor-evoked glomerular activity patterns

After demonstrating that sampling behavior does not underlie the differences in ODT between monomolecular odorants and their binary mixtures, we now attribute the increased ODT to mechanisms of neuronal processing in the olfactory system. Thus, we next tested the hypothesis that the number and pattern of activated glomerular units and their respective activation strengths correlate with ODT. We chose the Euclidean distance (ED) as a suitable parameter to represent the extent of similarity in odor-evoked activity maps, both with regard to the pattern as well as the strength of activity. Maps were visualized in anesthetized mice using intrinsic optical signals (IOS) evoked by the same set of odorants used for behavioral training (Figure 6a). Binary mixtures evoked more overlapping glomerular activity patterns compared to monomolecular odorants as measured by the ED for all odor pairs tested (Figure 6b, paired t test, p < 0.0005, n = 5 mice). Enantiomers, octanols and carvones evoked very similar activity patterns (paired t-test, p = 0.1, n = 5 mice) that reflected longer and similar ODT measurements we observed during the discrimination behavior. Correlating ED measurements with the corresponding ODT revealed a robust relationship between map similarity and discrimination time for all monomolecular odorants and binary mixtures. A strong negative correlation (Figure 6c, R^2^ = 0.96, ANOVA, p < 10^-12^) between ED and ODT suggests that the time to discriminate two odors is based on the degree of similarity of their glomerular activity maps, as measured by the ED.

**Figure 6.**
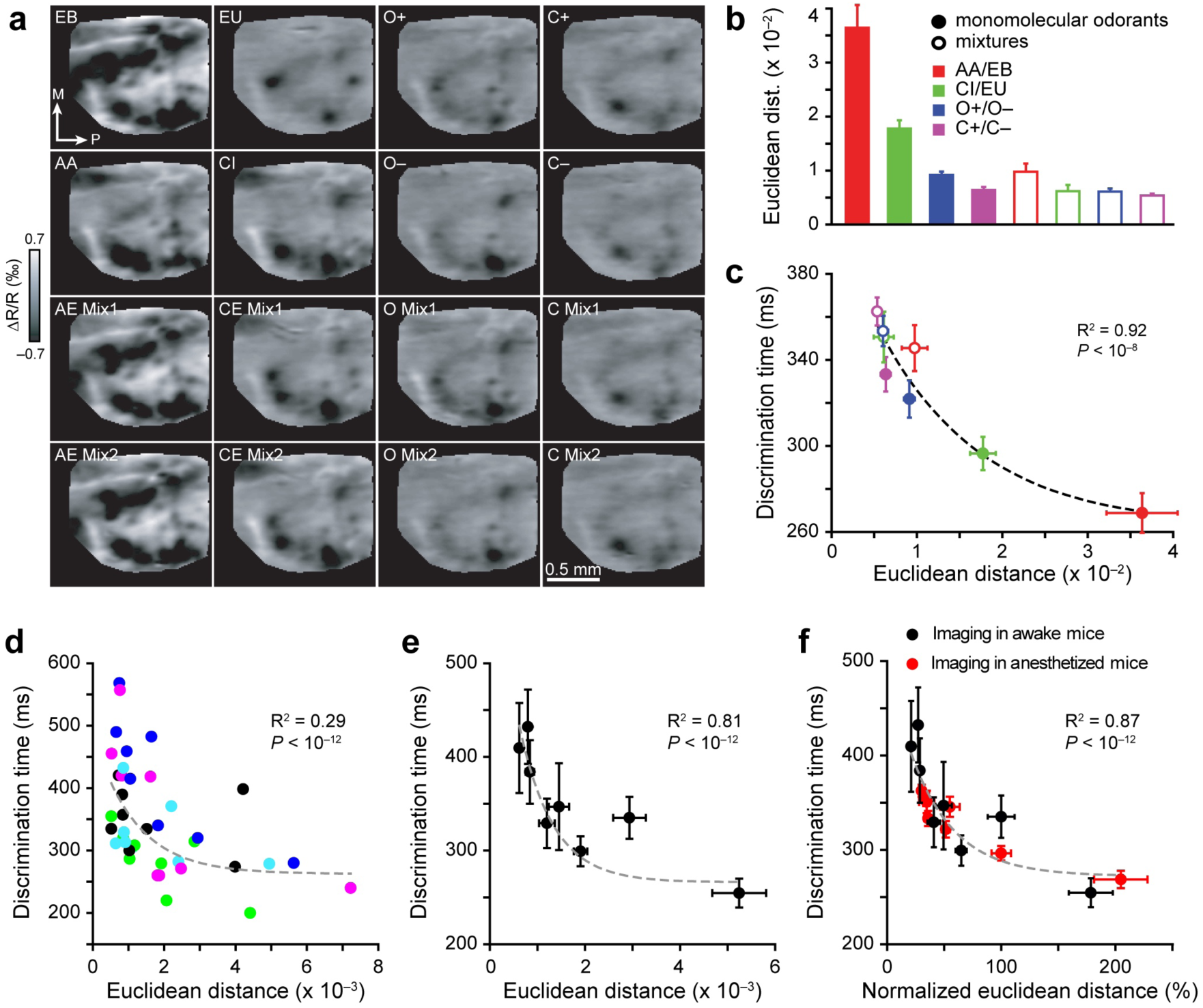
Euclidean distance between odor-activated glomerular maps in naïve and awake trained mice correlates with discrimination times measured for corresponding odor pairs. (a) IOS imaging of glomerular maps elicited by 4 pairs of monomolecular odors and their binary mixtures used in the behavioral training. (b) ED (see STAR Methods) between activation patterns evoked by different odor pairs. (c) Discrimination times measured for monomolecular odorants and binary mixtures are plotted as a function of ED between maps of activation patterns for corresponding odor pairs. Data were best fitted with a single exponential function (see STAR Methods). (d) ODTs measured during different discrimination tasks (10^-6^ IAA vs. 10^-6^ EB, 10^-4^ IAA vs. 10^-4^ EB, 10^-2^ IAA vs. 10^-2^ EB, 10^0^ IAA vs. 10^0^ EB, 10^-3^ CI vs. 10^-3^ EU, 10^-2^ Ci vs. 10^-2^ EU, 10^-1^ Ci vs. 10^-1^ EU, 10^0^ CI vs. 10^0^ EU) are plotted against the EDs measured for the corresponding odor pairs for individual mice. Dilutions were used to generate sparse representations in the OB. Colors indicate data collected for each individual mouse (n = 5). (e) ODTs and ED measured are averaged across mice (n = 5 mice). (f) ODTs measured for 16 odor pairs is plotted as a function of normalized ED, both from awake and anesthetized mice (normalized to 10^0^ Ci vs. 10^0^ Eu, which was common to both sets of experiments). Data are presented as mean ± SEM

### Euclidian distance predicts discrimination time in awake trained mice

Since ED measurements for monomolecular odorants and binary mixtures were done in untrained anesthetized mice, we further tested whether this correlation only holds for monomolecular-binary mixture comparison and if the discrimination training influences the correlation (Abraham et al., 2014). The latter is particularly important because the ODT is measured only after mice reached the learning criterion. Moreover, mice can detect and discriminate odorants even with the input from a single glomerulus (Smear et al., 2013). Therefore, we decided to test the discrimination abilities of mice over a wide range of concentrations, designed to activate different numbers of glomeruli, for two odor pairs where the lowest discriminable concentrations activated only 1-2 glomeruli on the dorsal surface. A batch of mice was trained on different dilutions of CI/EU and IAA/EB ranging from 10^0^ to 10^-10^ (percentile dilution in mineral oil). Mice started to learn the discrimination tasks from 10^-3^ for CI/EU and from 10^-6^ for IAA/EB towards lower dilutions. We measured the ODT for each dilution and correlated it with the ED measurements from imaging the same mice using an awake head-restrained setup. We observed the same negative correlation on single animal basis (Figure 6d, R^2^ = 0.29, ANOVA, p < 10^-12^) as well as when averaging across all mice used for imaging (Figure 6e, R^2^ = 0.81, ANOVA, p < 10^-12^). The absolute ED measurements varied in awake mice compared to the anesthetized situation due to different experimental conditions (see STAR Methods). To validate the ED measurements done under different conditions, we normalized the ED measurements to a common odor pair (CI vs. EU at 10^0^ dilution). We observed a negative correlation independently of the awake or anesthetized situation (Figure 6f, R^2^ = 0.75, p < 10^-12^). These results show that the map similarity measured by ED, reflecting both the number of activated glomeruli as well as their activation strength, can predict the time needed to discriminate an odor pair.

The ED between two vectors reflects not only the relative changes in the ratio of their values, but rather the difference in their absolute values, as opposed to the Pearson correlation coefficient. Therefore, we conclude that the properties of the glomerular patterns that are essential for odor discrimination are the identity of the activated glomeruli, their number and their activation strength. As these parameters define neuronal representations of the odorants, they can be employed as a neural metric for predicting olfactory discrimination time.

## Discussion

Here we show that the sniff-invariant decision time required by mice to discriminate odors is defined by the ED of their glomerular representations in the olfactory bulb. The degree of similarity of the spatial patterns as well as the number and strength of activated glomerular units define the ODT of a particular odor pair. The ODT is predicted to reach an upper limit of approximately 400-500 ms, when the ED between two glomerular activity patterns is minimized and further olfactory sampling may not improve discrimination performance. A time difference of up to 100-150 ms was found when comparing monomolecular odorants with the shortest and longest ODTs, or when comparing discrimination of binary mixtures with their monomolecular components. We propose that this difference in time reflects the time needed to refine (Abraham et al., 2010) and integrate information for complete percept formation (Chapuis and Wilson, 2012), facilitating accurate decisions in the discrimination process. Intriguingly, this processing time is independent of the sampling behavior used by the mice and accrues downstream of sampling. Our findings establish a neural metric of ODT and define the temporal windows of olfactory discrimination behavior as well as olfactory information processing in preparation for sniff-invariant decision-making.

### Behavioral paradigm

In the go/no-go olfactory behavior experiments with freely moving mice, ODTs were determined by the head-withdrawal behavior towards the unrewarded odors (Abraham et al., 2004), whereas in the head-restrained behavioral experiments, ODTs are determined from licking reactions towards the rewarded odors (Abraham et al., 2012). This comparison helps to control a possible confounding effect of motor behavior, because the latter may involve a less complex motor program than the former. However, our results suggest that the different motor responses and reward categories do not affect the measured ODT. As per the learning rules set for the head-restrained experiments, mice are supposed to register their first lick only within the first second of odor presentation, which allows them ample time to respond. Therefore, this design can detect the possible influence of urgency (for decision-making) on temporal correlates of sensory discrimination (Reddi and Carpenter, 2000).

### Sniff-invariant olfactory decisions

Rodents display different sniffing behaviors in a context-dependent manner (Wesson et al., 2008a), a strategy possibly designed to collect a maximal amount of information for better odor discrimination. Our breathing rate measurements show an increase in the sniff frequency specifically during the decision-making time period (Figures 3 and 4). This increase was invariant of chemical classes and difficulty of the tasks employed (Figure 5a) and it did not occur in the absence of odor stimuli (Figure S3). Thus, the time needed to discriminate odors of varying similarity is independent of the sampling behavior and puts our results in unison with the growing consensus that odor coding is sniff frequency-independent (Jordan et al., 2018a; Shusterman et al., 2018).

Animals show a transition from low frequency to high frequency sniffing when exposed to a novel odor source (Wesson et al., 2008a). The extent of such a shift depends on the behavioral context an animal is challenged with. When the animal is trained on a two-alternative choice task, allowing the animal to obtain reward for all stimuli on performing accurately, the increase in sniffing might be driven by reward anticipation (Kepecs et al., 2007). If the animals were discriminating two learned odorants, the observed increase in respiration frequency was almost 50% lower compared to an exploratory sniffing behavior (Wesson et al., 2008b). We observed an increase in the breathing frequency specifically during the decision-making period (Figure 3c) with a subsequent return to baseline. This is a clear indication that mice actively modulate their sniffing during the decision-making time window, yet independently of the difficulty of the discrimination task (Figure 5b).

### Determinants of the olfactory system governing discrimination time

The ODT includes time periods associated with air-flow to and across the olfactory epithelium, odor diffusion through mucus covering olfactory epithelium, binding of the odorant molecules to the olfactory receptor, signal transduction in olfactory sensory neurons (OSNs), olfactory processing in the OB and other olfactory areas, decision-making, initiation and programming of motor output, and execution of motor activity. At present it is difficult to attribute specific durations to each of these steps, but it is evident that differences in ODT could arise from odor-specific properties in any single or any combination of these steps. Recently, some studies investigated the role of sorptive properties of odorants in altering the odor representation and active sampling during the olfactory behavior (Cenier et al., 2013; Rojas-Libano and Kay, 2012). Although the physicochemical properties vary among the odorants we used, our results on sniffing prove that the ODT differences we do see between the monomolecular and binary mixtures is independent of animals’ sniffing behavior (Figure 5). Hence, studying sniffing behavior in conjunction with ODT allowed us to more specifically pin-point the time periods required for olfactory processing as illustrated in Figure 7. In this example of AA-EB binary mixture discrimination, the breath initiation counts (BICs) start to rise at 250 ms and peak at around 280-300 ms. These rising points vary for different odor pairs. Assuming that OSNs produce action potential output latest by the peak of the sampling curve (∼280 ms), the latency to the measured ODT (∼290 ms for AA/EB to ∼350 ms for C+/C- Mix, see Figure 5a) provides the time window needed to (1) complete odor representation in the OB and downstream processing centers, (2) decision-making and (3) motor initiation, programming and execution. If we consider the start point of the BIC rise, the dissimilar stimuli discriminations take ∼10-40 ms and similar stimuli discriminations need an additional 60-70 ms for the above mentioned processes under conditions of maximal performance (discrimination accuracy). While the exact time frame for each of these processes remains unknown, subtracting the two extremes yields the time exclusively consumed by processing in the OB and higher olfactory centers while discriminating highly similar stimuli. Based on our previous work (Abraham et al., 2010; Gschwend et al., 2015) we predict that the bulk of this time arises from the kinetics of inhibition in the OB. In turn, this notion predicts that processing in olfactory areas downstream of the OB occurs on very fast time scales. In conclusion, neural representations of olfactory information occur on a fast time scale ranging from 10 to 70 ms, consistent with previous work (Resulaj and Rinberg, 2015).

**Figure 7.**
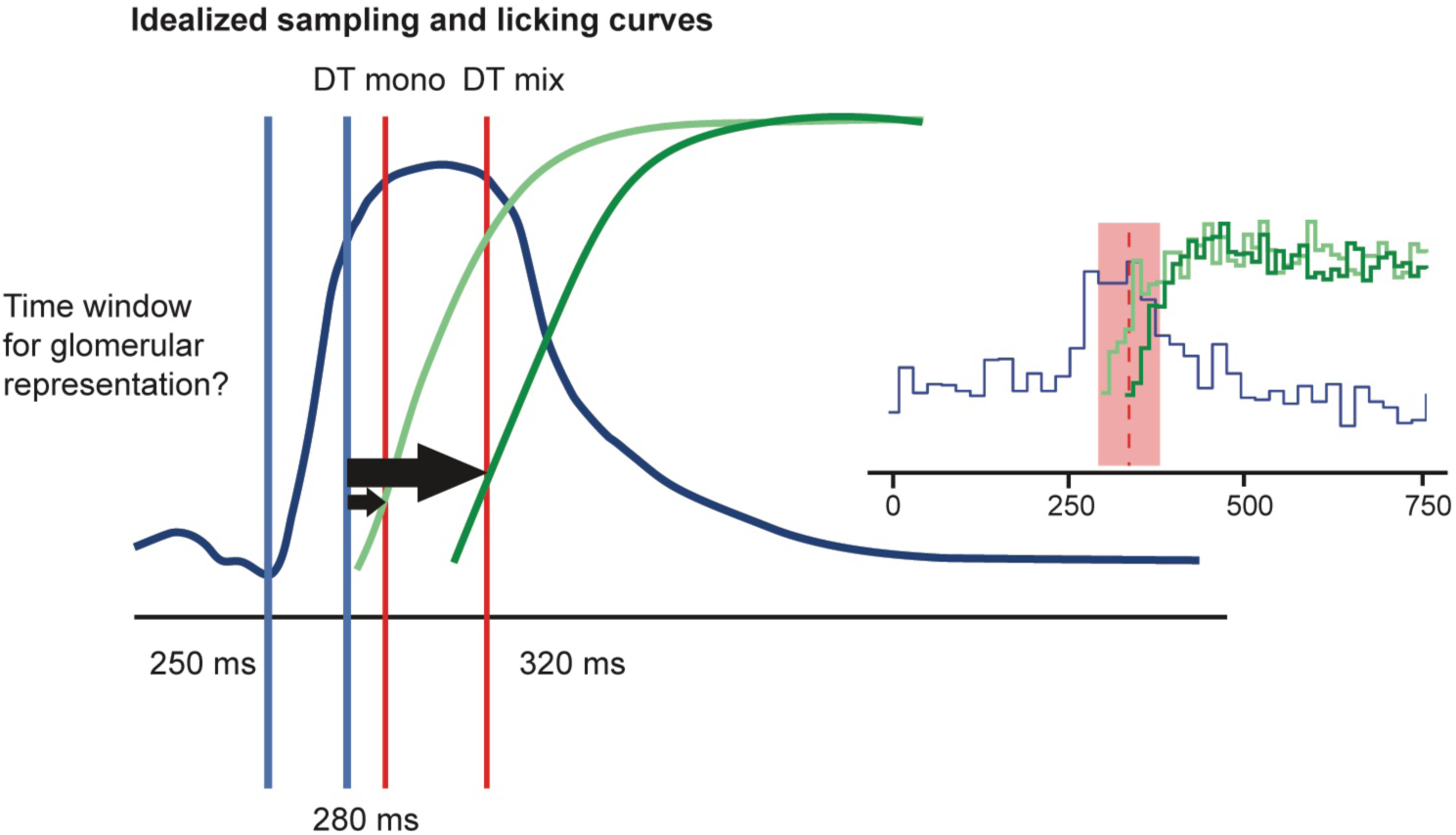
Fast odor discrimination supported by sniffing and licking behavior. An idealized sampling curve derived from the sniffing peak latency curve (blue curve) and the lick curves for simple and complex odors (light and dark green curves, respectively). The vertical blue lines delineate the brief phase of sniffing increase. The red vertical lines depict the ODTs for dissimilar and highly similar stimuli. The black arrows illustrate the times needed for representation and processing, assuming that odor has activated the olfactory sensory neurons latest at the time of peak sampling. The small black arrow illustrates a very brief decision-making time window for discriminating dissimilar stimuli (∼10 ms), while the large black arrow represents the much increased time window for discriminating highly similar stimuli (∼30 ms). Finally, the difference between the two arrows reflects only processing in the OB and downstream olfactory areas, because other contributions such as motor initiation, programming and execution should be cancelled out.

### A neural metric based on glomerular map similarity predicts odor discrimination time

For the following discussion, the similarity of two odorants is defined as the similarity of glomerular activation patterns elicited by the odorants, determined by IOS imaging of the dorsal OB in anesthetized and awake mice. Similarity was quantified by the ED between the two glomerular activity patterns, a parameter reflecting both geometrical distribution as well as number and activation strength of the glomerular units (small values of ED reflect high similarity). Each of the 16 odor pairs used in this study exhibited similarity-dependent differences in ODT within a range of up to 150 ms (Figure 6c,f). This holds true for discrimination of different pairs of monomolecular odorants as well as for binary mixture discrimination. Based on behavioral ODT measurements and *in vivo* imaging of glomerular activity patterns of 16 different odorant pairs, we propose a neural ‘olfactory’ metric according to which the ODT inversely scales with the ED derived from the glomerular activity patterns (Figure 6f).

To test if the olfactory metric applies also to sparse glomerular activation, we applied low concentrations of odorants only activating one or two glomeruli. Even with this sparse input, mice were able to learn the discrimination task and the measured ODTs correlated with the ED measurements (Figure 6d), strongly supporting the validity of the proposed neural olfactory metric.

### Application of the olfactory metric to discrimination of monomolecular odorants and their binary mixtures defines an upper limit of odor discrimination time

The ODTs of binary mixtures were increased in a manner anti-correlated to the ODT required to discriminate the two monomolecular components: the increase in ODTs for binary mixtures was larger for odorant pairs that evoked different glomerular maps and were discriminated rapidly (e.g. AA/EB), but smaller for those evoking highly similar maps and requiring more time (e.g. C+/C-) (Figure 2). This relation may appear counter-intuitive at first glance, but monomolecular odorants that already generate a highly similar map ‘consume’ a long ODT and the small increase in similarity of the derived binary mixture will produce only a small increase in ODT. Consequently, a maximal ODT of approximately 400-500 ms can be estimated for close to identical glomerular activity maps with small EDs (Figure 6c). Importantly, this figure is specific to the olfactometers used here and may deviate, depending on olfactometer design. A similar limiting value can be derived when comparing the increase of ODT for binary mixture discrimination relative to the ODT for discrimination of the monomolecular components (Figure 2b). Within the maximal ODT window, the olfactory system can produce highly accurate decisions exceeding 90% correct choices. In case of highly complex discriminations such as white odors (Weiss et al., 2012), the OB circuitry and downstream areas may not manage to separate and complete the patterns sufficiently well, even by consuming the maximal ODTs. In these situations, decision-making is based on incomplete percepts with the consequence of reduced accuracy. Mechanistically, the predicted upper limit of ODT may reflect a maximal processing time available to the neuronal mechanisms of pattern separation in the OB and downstream olfactory areas before a decision must be rendered.

### Stimulus-dependency of discrimination time

The stimulus-dependency of ODT remains an intensely discussed issue (Abraham et al., 2004, 2010, 2012; Ditzen et al., 2003; Frederick et al., 2011; Friedrich, 2006; Kepecs et al., 2006; Mainen, 2006; Rajan et al., 2006; Schaefer and Margrie, 2007; Uchida et al., 2006; Uchida and Mainen, 2003; Zariwala et al., 2013). The data reported here supports the idea of stimulus-dependency by generalizing this property, observed so far in the odorant pairs AA/EB (Abraham et al., 2004, 2010, 2012) and enantiomers of carvones (Rinberg et al., 2006), to additional odor pairs. With all four classes of stimuli tested, esters, CI/EU and two enantiomers, we observed a similarity-dependent ODT for the discrimination of monomolecular odorants as well as binary mixtures. Therefore, several lines of evidence strongly support stimulus-dependency in mice, consistent with other sensory systems (Luce, 1986).

However, behavioral tests in other species (honeybee and rat) revealed stimulus-independence (Ditzen et al., 2003; Uchida and Mainen, 2003; Zariwala et al., 2013), leaving the question unresolved at a larger scale. In case of the honeybee, differences in the design of the olfactory system due to the large phylogenetic distance may provide an explanation, but the similarity-independent ODTs observed in rats (Uchida and Mainen, 2003; Zariwala et al., 2013) are inconsistent with our conclusions. An article by Zariwala et al., challenged the classical concept of speed-accuracy tradeoff in decision-making (Zariwala et al., 2013). In their two-choice task, the stimulus is offered from a central port, and, depending on the stimulus, rats are forced to move either right or left to receive the reward. Odor categorization was achieved in a definite temporal window independent of task difficulty, although there was a difference of 30 ms between the least and most similar stimuli in certain conditions. Strikingly, in our paradigm we observed a similar time difference of approximately 30 ms between the monomolecular pair and binary mixtures of enantiomers, reflecting the time required for olfactory information processing (see discussion above). Moreover, the experiments reported by Zariwala et al., (2013) were based on binary mixtures of a single odor pair, octanols. The use of one odor pair, however, limits generalized conclusions on the existence of speed versus accuracy tradeoff and stimulus-dependency in olfaction. Our study overcomes this limit by demonstrating stimulus-dependency in more than one class of olfactory stimuli.

### Urgency, motivation and discrimination speed

Behavioral contingencies such as motivational status are important parameters that can influence the decision-making process (Gold and Shadlen, 2007). In our go/no-go freely moving paradigm, the ODT is based on the head-withdrawal behavior of mice towards non-rewarded odors (Abraham et al., 2004). The major advantage of this approach is that mice can afford an amount of time sufficient for accurate decisions, because they do not benefit from quick decisions. Therefore, the ‘urgency to react’ did not influence the decision-making time (Reddi and Carpenter, 2000) in the experiments reported herein. A possible disadvantage concerns a decrease in motivation coupled to the reaction towards the non-rewarded stimulus. To address this problem, we used a head-restrained behavioral paradigm (Abraham et al., 2012), where the ODTs are primarily based on the reactions towards a rewarded odor measured by licking behavior. In both cases animals performing at high accuracy took additional time for the discrimination of binary mixtures (Figures 2 & S1). Hence, the olfactory system shares the common feature of fast and similarity-dependent discrimination time observed in other sensory systems (Luce, 1986). This discrimination time can be predicted by the strength and similarity of the OB input patterns.

## Author Contributions

N.A., T.K., A.C., A.B. and S.K. carried out the study conceptualization and experimental design. N.A., A.B. S.K., D.N. and A.D. performed and analyzed behavioral experiments and sniff measurements. D.T. R.V. and H.S. acquired and analyzed intrinsic signal imaging data. N.A., T.K. and A.C. wrote the manuscript with comments from A.B.

## Acknowledgements

We thank Andreas Schaefer, Tansu Celikel, Raghav Rajan, N.K. Subhedar, Collins Assissi and Laboratory of Neural Circuits and Behaviour (LNCB) members for fruitful discussions. We thank staff of National Facility for Gene Function in Health and Disease (NFGFHD) for the technical support. This work was supported by the Wellcome Trust – DBT India Alliance intermediate grant (IA/I/14/1/501306 to N.A.), UGC NET Fellowship (A.B.), IISER-Pune Fellowship (S.K.), Heidelberg Academy of Sciences (H.S. and T.K.), DFG Research Unit grant FOR-643 (H.S. & T.K.), DFG Special Priority Programme grant SP1134 (T.K.), the European community (H2020-FETOPEN-2014-2015-662629-NanoSmell to A.C.), the Swiss National Science Foundation (grant 31003A_172878 to A.C.), Postdoc program of Medical Faculty of Heidelberg University (N.A.), and the European Molecular Biology Organization (N.A. long-term postdoctoral fellowship program).

## Star Methods

### Contact for Reagent and Resource Sharing

Further information and requests for reagents may be directed to, and will be fulfilled by the corresponding author Nixon M. Abraham.

### Experimental Model and Subject Details

#### Mice

C57BL/6J mice (Jackson Laboratories and Charles River) were used for the behavior and imaging experiments. Subjects were 5–6 weeks old at the beginning of the behavioral experiments and maintained on a 12 hr light-dark cycle in a temperature- and humidity-controlled animal facility. All behavioral training was conducted during the light cycle. During the training period, animals had free access to food but were on a water restriction schedule designed to keep them at >85% of their baseline body weight. Continuous water restriction was never longer than 12 hr. All animal care and procedures were in accordance with the Institutional Animal Ethics Committee (IAEC) at IISER Pune, the animal ethics guidelines of the Max Planck Society, the University of Heidelberg and the University of Geneva, Swiss Federal Act on Animal Protection and Swiss Animal Protection Ordinance and the Committee for the Purpose of Control and Supervision of Experiments on Animals (CPCSEA), Government of India.

### Method Details

#### Behavioral training under freely moving conditions

Olfactory discrimination experiments were performed using four modified eight-channel olfactometers (Knosys, Washington) controlled by custom software written in Igor (Wavemetrics, OR). Groups were usually counterbalanced between setups. In brief, odorant from one out of eight odor channels was presented to the mice in a combined odor sampling/reward port. This ensured tight association of the water reward with the presented odorant. Odorants were diluted in mineral oil (Fluka, percentage of dilution is indicated in figure legends) and further diluted 1:20 by airflow. Each rewarded odor (S+) and non-rewarded (S-) odor was presented from as many valves as possible (usually four each). Odors were made up freshly for each task (generally every day). Behavioral training consisted of a task habituation phase followed by discrimination training phase. Discrimination times were measured during the training phase while animals were performing with high accuracy.

##### Task habituation phase

Beginning 1-3 days after the start of the water restriction schedule, animals were trained using standard operant conditioning procedures (See also Abraham et al., 2004). In a first pre-training step, each lick at the water delivery tube was rewarded. After 20 licks a second stage began in which head insertion initiated a 2 s “odor” presentation during which a lick was rewarded. The “odorant” used in the pre-training was the mineral oil used for odor dilution. The complexity of the pre-training task increased gradually during five phases, each consisting of 20 trials. Essentially all animals learned this task within one day (2-3 sessions of 30 min each).

##### Discrimination training phase

The mouse initiates each trial by breaking an infrared beam at the sampling port opening [For details, see also (Abraham et al., 2004)]. This opens one of eight odor valves and a diversion valve (DV) that allow to divert all air flow away from the animal for a variable time (usually t_DV_ = 500 ms). The use of the diversion valve ensures that odor traveling time between “odor onset” and first contact of the animals’ nose with the odor is minimized. We thus do not correct for any estimated odor traveling time and present the raw, unedited discrimination times throughout the paper. After the release of the DV, the odor is applied to the animal for 2 s. If the mouse licks at least once in three 500 ms time bins out of four at the lick port during this time it can receive water reward of 2-4 µl at the end of the 2 s period. Trials were counted as correct if the animals met the criterion we set for water delivery (licking at least once in three out of four 500 ms bins) during the S+ presentation or if animal did not lick continuously for S-. A second trial cannot be initiated unless an ITI of at least 5 s has passed. This interval is sufficiently long so that animals typically retract quickly after the end of the trial. The minimal inter-stimulus interval is thus 5 s, which seemed to be sufficient as no habituation could be observed (performance was not correlated with the actual ITI chosen by the animal, which was around 10-20 s). No minimal sampling time is required to artificially enforce the animal to take a decision.

Odorants are presented in a pseudo-randomized scheme (no more than 2 successive presentations of the same odor, equal numbers within each block of 20 trials). No intrinsic preference towards any of the odors was observed. Bias by odor preferences was generally avoided by counterbalancing S+ and S-stimuli such that each odor was designated S+ or S-stimulus for the same number of animals. A total of 200-300 trials were performed each day, which are separated into 30-40 min stretches to ensure maximal motivation despite the mildness of the water restriction scheme. Motivation was controlled by monitoring ITI and the lick frequency. Animals that did not reliably insert their head into the odor port to initiate a trial were excluded from the analysis (∼5% of all animals). The animals that kept their head in the odor port for S-trials without licking were also excluded from the analysis (∼5% of all animals).

##### Measurement of discrimination times

The sampling behavior of mice is monitored by the status of the infrared beam inside the sampling chamber from where they sense the odor and receive the reward (For details, see also (Abraham et al., 2004)). Upon presentation of a S+ odor, the animal continuously breaks the beam, whereas upon presentation of an S-odor an animal familiar with the apparatus usually quickly retracts its head. The average difference in response to the S+ and S-odor is approximately sigmoidal and yields a sensitive measure of the discrimination performance. Reaction times were determined as follows: Combining 300 successive trials, for every time point, beam breaking for S+ and S-odors were monitored in the order of microseconds and finally pooled as 20 ms time bins and then compared, yielding significance value as a function of time after odor onset. The last crossing of the p = 0.05 line was measured by linear interpolation in the logarithmic plot and determined the discrimination time.

##### Measurement of olfactory discrimination times for different odor pairs

The first set of experiments consisted of 10 male mice (set 1), the second set consisted of 7 mice (set 2) and third set consisted of 6 mice (set 3). In each of the blocks ODT was measured from the last 300 trials.

The following sequence of odorants was applied in set 1 (Figures 1a-d and 2a-b, numbers reflect the number of trials): 1200 CI vs. EU, 600 AA vs. EB, 900 AA60/EB40 vs. AA40/EB60, 600 AA vs. EB, 900 AA60/EB40 vs. AA40/EB60, 900 AA54/EB46 vs. AA46/EB54 (results not reported here, axis break in Figure 1), 600 CI vs. EU, 900 CI60/EU40 vs. CI40/EU60, 600 CI vs. EU, 900 CI60/EU40 vs. CI40/EU60, 900 CI80/EU20 vs. CI20/EU80 (results not reported here, axis break in figure 1), 900 C+ vs. C– (9 mice, one mouse had to be dropped due to lack of motivation), 900 C+/C– 60/40 mix (9 mice), 600 C+ vs. C–, 900 C+/C– 60/40 mix.

The following sequence of odorants was applied in set 2 (Figures 1e-h and 2a-b): 1200 CI vs. EU (not reported), 900 O+ vs. O–, 900 O+/O– mix, 600 O+/O–, 900 O+/O– mix (7 mice). Odorants were presented at 0.1% dilution in MO.

The third set of mice (n = 6) were trained on CI vs. EU (1200 trials), followed by octanols (900 trials) and their binary mixtures (1200 trials). Odorants were presented at 1% dilution in MO. Performance and ODT were not different from the ones reported for set 2 (data pooled in Figure 2).

Fourth batch of mice were trained under head-restrained conditions (see next section) on the above-mentioned odor pairs for investigating the sniffing behavior while animals were actively involved in the olfactory behavioral training (n = 7-8 mice, set 4). To study the sampling behavior in the absence of any odors, we trained another batch of naïve mice MO vs. MO (Figure S2, n = 8 mice, set 5).

To study the glomerular representation of the odorants used for the behavioral training, a batch of naïve mice (n = 5 mice, set 6, see later) were used for the intrinsic optical signal imaging. The seventh set of mice (n = 5) were trained on different dilutions of CI/EU and IAA/EB ranging from 10^0^ to 10^-10^ (percentile dilution in mineral oil). These mice were used for imaging under awake condition (Figure 6).

#### Behavioral training under head-restrained conditions

The experimental protocol is detailed in Abraham et al., 2012. In brief, before each behavioral session, a mouse was placed in a PVC tube and head-restrained by screwing the head post on a custom made metallic device fixed on a platform. All olfactory discrimination behavior experiments were done using a custom built olfactometer, which was synchronized to a licking circuit and breath measuring device (lickometer) for recording and digitizing the lick and breath responses of mice towards different odorants (Abraham et al., 2012; Bathellier et al., 2007, 2008; Gschwend et al., 2012, 2015). Breathing pattern of animals were recorded using an airflow pressure sensor (Figure S2) while animals were actively involved in olfactory behavior experiments. Mice were trained for the following sequence of odorants: CI vs. EU, CI60/EU40 vs. CI40/EU60, AA vs. EB, AA60/EB40 vs. AA40/EB60, C+ vs. C–, C+/C– 60/40 mix, O+ vs. O–, O+/O– 60/40 mix (n = 7-8 mice, set 4). To calculate the basal breathing parameters a fifth batch of naïve mice were trained for MO vs. MO (Figure S3, n = 8 mice, set 5).

Odor mixing and dilutions were achieved by the olfactometer through which a clean stream of air was split into a dilution stream and an odor stream that passed through bottles containing saturated odorants. These streams were merged right before the output at the level of an odor delivery nozzle. The settings of olfactometer allowed us to obtain a sharp stimulation onset and concentration stability during odor presentation (Patterson et al., 2013). Odor concentration is expressed as a percentage of the saturated vapor pressure, which reflects the relative flow rates of the odor and dilution streams. For example, a value of 5% means that the saturated odor stream was set at a flow rate 20 times lower than the dilution stream. In all cases, the total output flow rate was equal to 400 sccm. Odorants were diluted to 1% in mineral oil and further diluted by airflow. Odors were freshly prepared for each task. Behavioral training under head-restrained conditions started with a habituation training followed by discrimination training phase. Discrimination times were measured while animals performed with high accuracy. Sniffing was recorded throughout the training phase.

##### Task habituation training

Beginning 1-3 days after starting the water restriction schedule, 8 mice (set 4) were trained for an associative task using operant conditioning procedures. In a first stage (20 trials), immediately after a tone of 200 ms (5000-6000 Hz), a water drop (2 µl) was presented to animals irrespective of their responses. In the subsequent trials the delay between the tone and the water reward was increased to 1 s (20 trials) and 2 s (20 trials). This was meant to provoke the licking behavior by animals. Once the animals learned to wait for the water reward, we introduced an odor of mineral oil (the solvent used to dilute the odorant) for 2 s before the water reward. During a second stage, following the tone, a baseline recording of licking (1 s) was performed before 2 s presentation of 1% methyl benzoate. If animals were not licking during the baseline, we implemented the criterion for water delivery based on their licking time during the odor presentation. The total licking time required, during odor presentation, to trigger water reward was gradually increased in each step from 40 ms up to 240 ms (40 ms – 30 trials, 80 ms – 30 trials, 120 ms – 30 trials, 160 ms – 50 trials, 240 ms – 50 trials). If animals were licking during the baseline, the required licking time kept increasing from 100% (same amount of licking as during the baseline) up to 200% (100% – 30 trials, 125% – 30 trials, 150% – 30 trials, 175% – 50 trials, 200% – 50 trials). Essentially most animals learned this task in 2-3 days (4-6 sessions of 30 min each).

##### Olfactory behavioral training

Trial initiation was set by the experimenter with a constant ITI (13.2 s) between consecutive trials for all mice used in the experiment. For an odor discrimination task, mice were trained for two odorants, one being rewarded (S+) and the other being unrewarded (S-). The required total licking duration for getting the water reward (2 µl) at the end of a rewarded trial was based on the licking activity during baseline. After the recording of baseline licking for 1 s, the odor was applied to the animal for 2 s. If mice licked during the baseline, they had to lick double amount of time during the odor presentation to get water reward. If mice were not licking during the baseline, the criterion for a water reward was a total lick time of 80 ms in three time bins of 500 ms out of four bins during the 2 s odor presentation. Trials were counted as correct if the animal met with the above-mentioned criteria for rewarded trials. For unrewarded trials, if mice were not licking during baseline, the criterion for a correct trial was a maximum lick time of 80 ms in one time bin of 500 ms out of four during the 2 s odor presentation. If mice licked during the baseline of an unrewarded trial, the trial was counted as correct if the total licking time during 2 s odor presentation did not exceed 25% of their baseline licking. Generally, most of the mice did not lick for unrewarded trials and they consistently licked for rewarded trials after the task acquisition. No punishment was given to the mice for incorrect trials.

Odors were presented on a pseudo-randomized scheme (no more than 2 successive presentations of the same odor, equal numbers within each block of 20 trials, ensuring different order of presentations for S+ and S-trials within each 20 trial blocks). No intrinsic preference towards any of the odors was observed. Bias caused by odor preferences was generally avoided by counterbalancing S+ and S-stimuli such that each odor was designated S+ or S-stimulus for the same number of animals. A total of 200-300 trials per animal, separated into 30-40 min sessions to ensure maximal motivation despite the mildness of the water restriction scheme, were performed each day. Motivation was controlled by the frequency of licking. When the animals were unmotivated, they stopped responding to the rewarded trials.

##### Measurement of discrimination time

The licking behavior of mice was monitored with a high temporal resolution of 2 ms. Upon presentation of a S+ odor, the animals continuously licked during the odor, whereas upon presentation of an S-odor mice hardly licked during the odor presentation. The average difference in response to the S+ and S-odor is approximately sigmoidal and yields a sensitive measure of the discrimination performance. Reaction times were determined as follows: The tasks with ≥80% performance levels were selected and combined for 300 successive trials [(150 S+ and 150 S-), 20 trial-blocks with low accuracy levels were excluded for DT measurements]. For every 2 ms time point licking was monitored and then compared between S+ and S-, yielding significance value as a function of time after odor onset. The last crossing of the p = 0.05 line was measured by linear interpolation in the logarithmic plot and determined the discrimination time.

#### Measurement of breathing parameters

The breathing patterns of mice during behavioral training were continuously recorded with an airflow pressure sensor (Honeywell AWM2300V) placed near one of the nostrils. The breathing circuit returns a digital signal in terms of raw voltage at a temporal resolution of 4 µs. Unfiltered traces were recorded with Tektronix TBS 1072B-EDU oscilloscope using a custom-written code in Python. Traces were triggered by a tone marking the beginning of trials and were recorded for 10 seconds. A threshold function was applied to the voltage signal to give a binary signal where a value of 0 is equivalent to low voltage (inhalation) and a value of 1 is equivalent to high voltage (exhalation, Figure S2). The filtered breathing traces were collected continuously using the custom-written acquisition software.

##### Calculation of breathing frequencies and histograms

The inspiration marks the beginning of a breathing cycle. For each trial in the behavioral task, the breathing frequency was calculated for the entire stimulus period (Figure 3a, b). For each odor pair, blocks of 20 trials with ≥ 80% performance levels were selected and combined for 300 successive trials (150 S+ and 150 S-). All time-points at which a breathing cycle started were marked in a raster plot (Figure 4, lower part of panels). The distribution of breathing cycles over all trials was visualized in a histogram using a bin size of 20 ms. (Figure S2). The time at which this histogram peaks (mode of the distribution of breathing cycles) was defined as the sniffing peak.

#### *In vivo* optical imaging

For imaging in anesthetized animals (n = 5, set 6), mice were anesthetized using urethane (1.5 g/kg i.p.). Heart and respiration rate were continuously monitored. Anesthetic was supplemented throughout the experiments as needed. The body temperature was kept between 36.5°C and 38°C using a heating pad and a rectal probe (FHC, Bowdoinham, ME). For imaging in awake animals (n = 5, set 7), preparations and head-post implantations, necessary to perform experiment in head-restrained conditions, were done as described previously (Vincis et al., 2012). Prior to imaging sessions, each mouse was habituated to head-restrained condition for 2-4 sessions (30 min each), over 2-3 days.

The OB surface was illuminated using a light guide system with a 700 ± 15 nm interference filter and a halogen lamp as light source. At the beginning and the end of each experimental session, images of the blood vessel pattern were taken using green light (546 nm interference filter) in order to assess the focus and to minimize potential drift. Two different camera systems were used for imaging. The first comprised a macroscope (Navitar 17; N.A. 0.46; Nikon 135 mm; f=2.0) and the Imager 3001F optical system (Optical Imaging, Mountainside, NJ); the second consisted of a modified Fuji camera (HR Deltaron 1700), the Imager 2001VSD+ controller and the DyeDaq software package. Images were acquired at either 5 Hz during 10 s or at 6.5 Hz during 9.3 s. After a short baseline period without odor stimulus (2 or 3.5 s), odors were presented for several seconds (4-5 s).

Odorants were presented using custom-made olfactometers with individual odor lines and nozzles for each odor (see section “Behavioral training under under head-restrained conditions”) or a GC PAL robot injection system (CTC Analytics, Zwingen, Switzerland). In the latter case, odorant concentrations were controlled by adjusting the airflow rates by mass flow controllers. One part of the air stream (flow rate 25 ml/min, 50 ml/min, 100 ml/min or 200 ml/min) was led through air-saturator bottles containing 5 ml of pure (undiluted) odors. Subsequently, this air was mixed with a stream of clean air to achieve a final airflow of 2 l/min. The GC PAL robot injection system offered the possibility of a completely automated presentation of many odors within a short time. For this purpose, 400 µl of undiluted odors (monomolecular odors or odor mixtures) were pipetted into 20 ml vials. Vials were placed into sample trays. 2.5 ml of the odor-enriched vial head-space were transferred by a syringe and injected into a stream of clean air (flow rate: 2 l/min). This air stream was led through a Teflon tube to the stimulator, which was placed 1-2 cm in front of the nose of the animal. After each odor presentation, the syringe was flushed with nitrogen for 30 seconds. Inter-stimuli intervals were at least 60 seconds long to minimize adaptation and desensitization effects. The order of stimuli presentation was varied among experiments; in all cases, no influence of the order of stimuli presentation on the outcome of the analysis could be observed.

Data analysis was performed using custom-written scripts in the Matlab programming environment (The Mathworks, Natick, MA). All frames were divided by the baseline reflectance (the “first frames”) for normalization. A two-dimensional Gaussian band-pass filter (σ low-pass = 13 μm, σ high-pass = 200 μm) was applied to remove global nonspecific signals and high-frequency noise. A number of frames (the “active frames”) were averaged for each odorant presentation to obtain one single activity map. Although single odor presentations were sufficient to obtain distinct functional maps, usually four to six odor presentations were averaged to improve the signal-to-noise ratio. Regions beyond the bulb surface and regions producing artifacts (big blood vessels and pigment cells) were masked using one exclusion mask for each animal. The masked regions were not used for map comparison. Each map (a digital black-and-white image) was represented as a vector of values. Each vector element represented a pixel value, as it was obtained by recording and subsequent data analysis (normalization and filtering, see above). Vector pairs were compared by calculating the Pearson correlation and the ED between them. For the calculation of the ED a vector can be thought of as a point in a multidimensional space, with the number of vector elements corresponding to the number of dimensions. In each dimension, the coordinate of the point corresponds to the value of the vector element representing this dimension. The distance between two points in the Euclidean space can be calculated using a simple formula (the Pythagorean formula). Normalization between animals was not necessary, since maps were always compared only within the same animal. However, for direct comparison between awake and anesthetized datasets (Figure 6f), we used the Euclidean distance calculated for the common odor pair (CI/EU) as a normalization factor for all other odorants since imaging equipment were different (optics and camera specification) and movement artifacts were more prominent in awake mice compared to anesthetized mice.

#### Quantification and Statistical analysis

In this study, statistical analyses were performed using Statistica, GraphPad Prism and OriginPro. We used ANOVA and associated post-hoc tests, two tailed paired t-test and different non-parametric tests (see text and legends). Shapiro-Wilk test was used to assess normality of the data. For all parametric ANOVA, homogeneity of variance was tested using Levene’s test or a test of sphericity (for one-way repeated measures ANOVA). Relationship between Euclidean distance and ODT (Figures 6c-f) were best fitted with a single exponential (equation: y = A1*exp(-x/t1) + y0, OriginPro).

## References

Abraham, N.M., Egger, V., Shimshek, D.R., Renden, R., Fukunaga, I., Sprengel, R., Seeburg, P.H., Klugmann, M., Margrie, T.W., Schaefer, A.T., and Kuner, T. (2010). Synaptic inhibition in the olfactory bulb accelerates odor discrimination in mice. Neuron 65, 399–411.

Abraham, N.M., Guerin, D., Bhaukaurally, K., and Carleton, A. (2012). Similar odor discrimination behavior in head-restrained and freely moving mice. PLoS One 7, e51789.

Abraham, N.M., Spors, H., Carleton, A., Margrie, T.W., Kuner, T., and Schaefer, A.T. (2004). Maintaining accuracy at the expense of speed: stimulus similarity defines odor discrimination time in mice. Neuron 44, 865–876.

Abraham, N.M., Vincis, R., Lagier, S., Rodriguez, I., and Carleton, A. (2014). Long term functional plasticity of sensory inputs mediated by olfactory learning. eLife 3, e02109.

Bathellier, B., Buhl, D.L., Accolla, R., and Carleton, A. (2008). Dynamic ensemble odor coding in the mammalian olfactory bulb: Sensory information at different timescales. Neuron 57, 586–598.

Bathellier, B., Van De Ville, D., Blu, T., Unser, M., and Carleton, A. (2007). Wavelet-based multi-resolution statistics for optical imaging signals: Application to automated detection of odour activated glomeruli in the mouse olfactory bulb. NeuroImage 34, 1020–1035.

Cenier, T., McGann, J.P., Tsuno, Y., Verhagen, J.V., and Wachowiak, M. (2013). Testing the sorption hypothesis in olfaction: a limited role for sniff strength in shaping primary odor representations during behavior. J Neurosci 33, 79–92.

Chapuis, J., and Wilson, D.A. (2012). Bidirectional plasticity of cortical pattern recognition and behavioral sensory acuity. Nature neuroscience 15, 155–161.

Ditzen, M., Evers, J.F., and Galizia, C.G. (2003). Odor similarity does not influence the time needed for odor processing. Chemical senses 28, 781–789.

Fletcher, M.L. (2011). Analytical processing of binary mixture information by olfactory bulb glomeruli. PLoS One 6, e29360.

Frederick, D.E., Rojas-Libano, D., Scott, M., and Kay, L.M. (2011). Rat Behavior in Go/No-Go and Two-Alternative Choice Odor Discrimination: Differences and Similarities. Behavioral neuroscience 125, 588–603.

Friedrich, R.W. (2006). Mechanisms of odor discrimination: neurophysiological and behavioral approaches. Trends in neurosciences 29, 40–47.

Friedrich, R.W., and Korsching, S.I. (1998). Chemotopic, combinatorial, and noncombinatorial odorant representations in the olfactory bulb revealed using a voltage-sensitive axon tracer. Journal of Neuroscience 18, 9977–9988.

Gold, J.I., and Shadlen, M.N. (2007). The neural basis of decision making. Annual review of neuroscience 30, 535–574.

Gschwend, O., Abraham, N.M., Lagier, S., Begnaud, F., Rodriguez, I., and Carleton, A. (2015). Neuronal pattern separation in the olfactory bulb improves odor discrimination learning. Nature neuroscience.

Gschwend, O., Beroud, J., and Carleton, A. (2012). Encoding odorant identity by spiking packets of rate-invariant neurons in awake mice. PLoS One 7, e30155.

Huston, S.J., Stopfer, M., Cassenaer, S., Aldworth, Z.N., and Laurent, G. (2015). Neural Encoding of Odors during Active Sampling and in Turbulent Plumes. Neuron 88, 403–418.

Johnson, B.A., Ho, S.L., Xu, Z., Yihan, J.S., Yip, S., Hingco, E.E., and Leon, M. (2002). Functional mapping of the rat olfactory bulb using diverse odorants reveals modular responses to functional groups and hydrocarbon structural features. Journal of Comparative Neurology 449, 180–194.

Jordan, R., Fukunaga, I., Kollo, M., and Schaefer, A.T. (2018a). Active Sampling State Dynamically Enhances Olfactory Bulb Odor Representation. Neuron.

Jordan, R., Kollo, M., and Schaefer, A.T. (2018b). Sniffing Fast: Paradoxical Effects on Odor Concentration Discrimination at the Levels of Olfactory Bulb Output and Behavior. eNeuro 5.

Kauer, J.S. (1988). Real-time imaging of evoked activity in local circuits of the salamander olfactory bulb. Nature 331, 166–168.

Kauer, J.S., and White, J. (2001). Imaging and coding in the olfactory system. Annual review of neuroscience 24, 963–979.

Kay, L.M., Beshel, J., and Martin, C. (2006). When good enough is best. Neuron 51, 277–278.

Kepecs, A., Uchida, N., and Mainen, Z.F. (2006). The sniff as a unit of olfactory processing. Chemical senses 31, 167–179.

Kepecs, A., Uchida, N., and Mainen, Z.F. (2007). Rapid and precise control of sniffing during olfactory discrimination in rats. Journal of neurophysiology 98, 205–213.

Koehl, M.A., Koseff, J.R., Crimaldi, J.P., McCay, M.G., Cooper, T., Wiley, M.B., and Moore, P.A. (2001). Lobster sniffing: antennule design and hydrodynamic filtering of information in an odor plume. Science 294, 1948–1951.

Luce, R. (1986). Response Times. Oxford Psychology Series No 8 Oxford University Press.

Ma, M., and Shepherd, G.M. (2000). Functional mosaic organization of mouse olfactory receptor neurons. Proceedings of the National Academy of Sciences of the United States of America 97, 12869–12874.

Mainen, Z.F. (2006). Behavioral analysis of olfactory coding and computation in rodents. Current opinion in neurobiology 16, 429–434.

Mori, K., Nagao, H., and Yoshihara, Y. (1999). The olfactory bulb: coding and processing of odor molecule information. Science 286, 711–715.

Patterson, M.A., Lagier, S., and Carleton, A. (2013). Odor representations in the olfactory bulb evolve after the first breath and persist as an odor afterimage. Proceedings of the National Academy of Sciences of the United States of America 110, E3340–E3349.

Rajan, R., Clement, J.P., and Bhalla, U.S. (2006). Rats smell in stereo. Science 311, 666–670.

Reddi, B.A., and Carpenter, R.H. (2000). The influence of urgency on decision time. Nature neuroscience 3, 827–830.

Ressler, K.J., Sullivan, S.L., and Buck, L.B. (1994). Information coding in the olfactory system: evidence for a stereotyped and highly organized epitope map in the olfactory bulb. Cell 79, 1245–1255.

Resulaj, A., and Rinberg, D. (2015). Novel Behavioral Paradigm Reveals Lower Temporal Limits on Mouse Olfactory Decisions. J Neurosci 35, 11667–11673.

Rinberg, D., Koulakov, A., and Gelperin, A. (2006). Speed-accuracy tradeoff in olfaction. Neuron 51, 351–358.

Rojas-Libano, D., and Kay, L.M. (2012). Interplay between Sniffing and Odorant Sorptive Properties in the Rat. Journal of Neuroscience 32, 15577–15589.

Rubin, B.D., and Katz, L.C. (1999). Optical imaging of odorant representations in the mammalian olfactory bulb. Neuron 23, 499–511.

Rubin, B.D., and Katz, L.C. (2001). Spatial coding of enantiomers in the rat olfactory bulb. Nature neuroscience 4, 355–356.

Schaefer, A.T., and Margrie, T.W. (2007). Spatiotemporal representations in the olfactory system. Trends in neurosciences 30, 92–100.

Shusterman, R., Sirotin, Y.B., Smear, M.C., Ahmadian, Y., and Rinberg, D. (2018). Sniff Invariant Odor Coding. eNeuro 5.

Smear, M., Resulaj, A., Zhang, J., Bozza, T., and Rinberg, D. (2013). Multiple perceptible signals from a single olfactory glomerulus. Nature neuroscience 16, 1687–1691.

Spors, H., and Grinvald, A. (2002). Spatio-temporal dynamics of odor representations in the mammalian olfactory bulb. Neuron 34, 301–315.

Spors, H., Wachowiak, M., Cohen, L.B., and Friedrich, R.W. (2006). Temporal dynamics and latency patterns of receptor neuron input to the olfactory bulb. J Neurosci 26, 1247–1259.

Uchida, N., Kepecs, A., and Mainen, Z.F. (2006). Seeing at a glance, smelling in a whiff: rapid forms of perceptual decision making. Nature reviews 7, 485–491.

Uchida, N., and Mainen, Z.F. (2003). Speed and accuracy of olfactory discrimination in the rat. Nature neuroscience 6, 1224–1229.

Uchida, N., Takahashi, Y.K., Tanifuji, M., and Mori, K. (2000). Odor maps in the mammalian olfactory bulb: domain organization and odorant structural features. Nature neuroscience 3, 1035–1043.

Vincis, R., Gschwend, O., Bhaukaurally, K., Beroud, J., and Carleton, A. (2012). Dense representation of natural odorants in the mouse olfactory bulb. Nature neuroscience 15, 537–539.

Wachowiak, M., and Cohen, L.B. (2001). Representation of odorants by receptor neuron input to the mouse olfactory bulb. Neuron 32, 723–735.

Weiss, T., Snitz, K., Yablonka, A., Khan, R.M., Gafsou, D., Schneidman, E., and Sobel, N. (2012). Perceptual convergence of multi-component mixtures in olfaction implies an olfactory white. Proceedings of the National Academy of Sciences of the United States of America 109, 19959–19964.

Wesson, D.W. (2013). Sniffing behavior communicates social hierarchy. Curr Biol 23, 575–580.

Wesson, D.W., Carey, R.M., Verhagen, J.V., and Wachowiak, M. (2008a). Rapid encoding and perception of novel odors in the rat. PLoS biology 6, e82.

Wesson, D.W., Donahou, T.N., Johnson, M.O., and Wachowiak, M. (2008b). Sniffing behavior of mice during performance in odor-guided tasks. Chemical senses 33, 581–596.

Xu, F., Liu, N., Kida, I., Rothman, D.L., Hyder, F., and Shepherd, G.M. (2003). Odor maps of aldehydes and esters revealed by functional MRI in the glomerular layer of the mouse olfactory bulb. Proceedings of the National Academy of Sciences of the United States of America 100, 11029–11034.

Zariwala, H.A., Kepecs, A., Uchida, N., Hirokawa, J., and Mainen, Z.F. (2013). The limits of deliberation in a perceptual decision task. Neuron 78, 339–351.

