## Supplemental Figures for "Similarity and strength of glomerular odor representations define neural metric of sniff-invariant discrimination time"

### Supplemental figures and legends

**a**

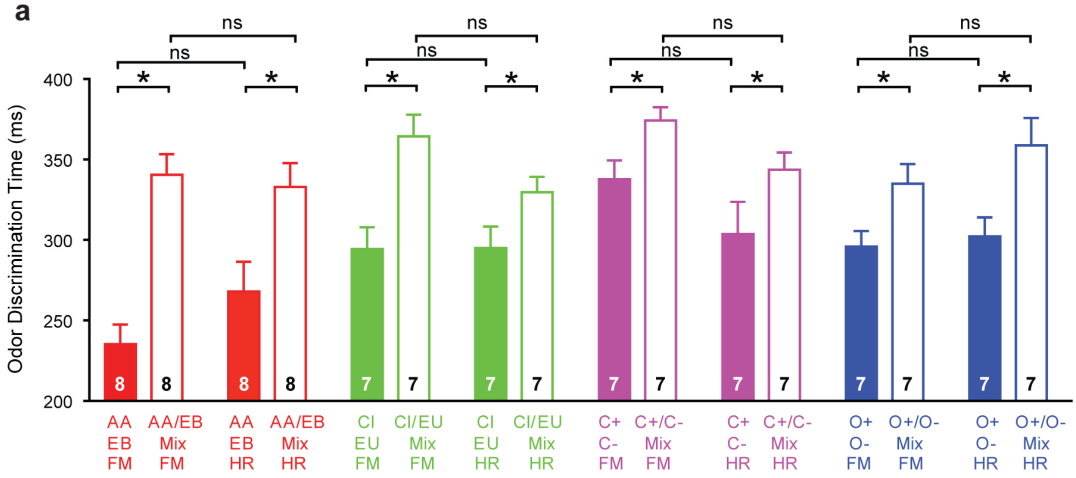


**Figure S1. Comparison of ODTs for simple monomolecular odorants and complex binary mixtures measured under freely moving and head-restrained experimental conditions, Related to Figure 2**

(a) Mice showed increased ODTs for the binary mixtures compared to the corresponding simple odors for all odor pairs tested under freely moving and head restrained conditions. Average ODTs across different experiments. Data are presented as mean ± SEM. $DT_{AA/EB(FM)}$ = 235 ± 14 ms, $DT_{AA/EBmix(FM)}$ = 341 ± 14 ms, $DT_{AA/EB(HR)}$ = 268 ± 19 ms, $DT_{AA/EBmix(HR)}$ = 333 ± 15 ms, $DT_{CI/EU(FM)}$ = 294 ± 14 ms, $DT_{CI/EUmix(FM)}$ = 365 ± 13 ms, $DT_{CI/EU(HR)}$ = 295 ± 13 ms, $DT_{CI/EUmix(HR)}$ = 330 ± 10 ms, $DT_{C+/C-(FM)}$ = 338 ± 13 ms, $DT_{C+/C-mix(FM)}$ = 374 ± 9 ms, $DT_{C+/C-(HR)}$ = 304 ± 20 ms, $DT_{C+/C-mix(HR)}$ = 344 ± 11 ms, $DT_{O+/O-(FM)}$ = 296 ± 10 ms, $DT_{O+/O-mix(FM)}$ = 335 ± 12 ms, $DT_{O+/O-(HR)}$ = 302 ± 12 ms, $DT_{O+/O-mix(HR)}$ = 359 ± 17 ms. Number of animals is indicated on each bar. * Comparison of ODTs for monomolecular odorants versus corresponding binary mixtures: Paired t test, $p < 0.05$)

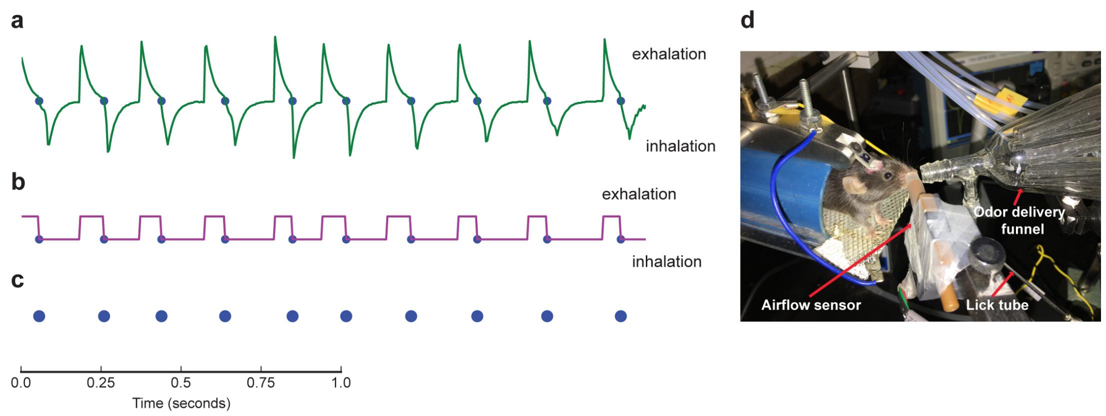


**Figure S2. Non-invasive measurement of breathing rate and derivation of breath initiation time points, Related to Figure 3**

(a) The breathing patterns of mice during behavioral training were acquired using an airflow pressure sensor placed near one nostril of the animal. Sensor voltage outputs are connected to the lickometer and displayed on an oscilloscope for live visualization of the breathing patterns.

(b) A threshold function is applied to the voltage signal to generate a binary signal where a value of 0 is equivalent to low voltage (inhalation) and a value of 1 is equivalent to high voltage (exhalation).

(c) Breath initiations are marked as blue dots. For each trial, the blue dots from the trial period are used to generate the SPL raster plots.

(d) Animal engaged in olfactory behavior experiment.

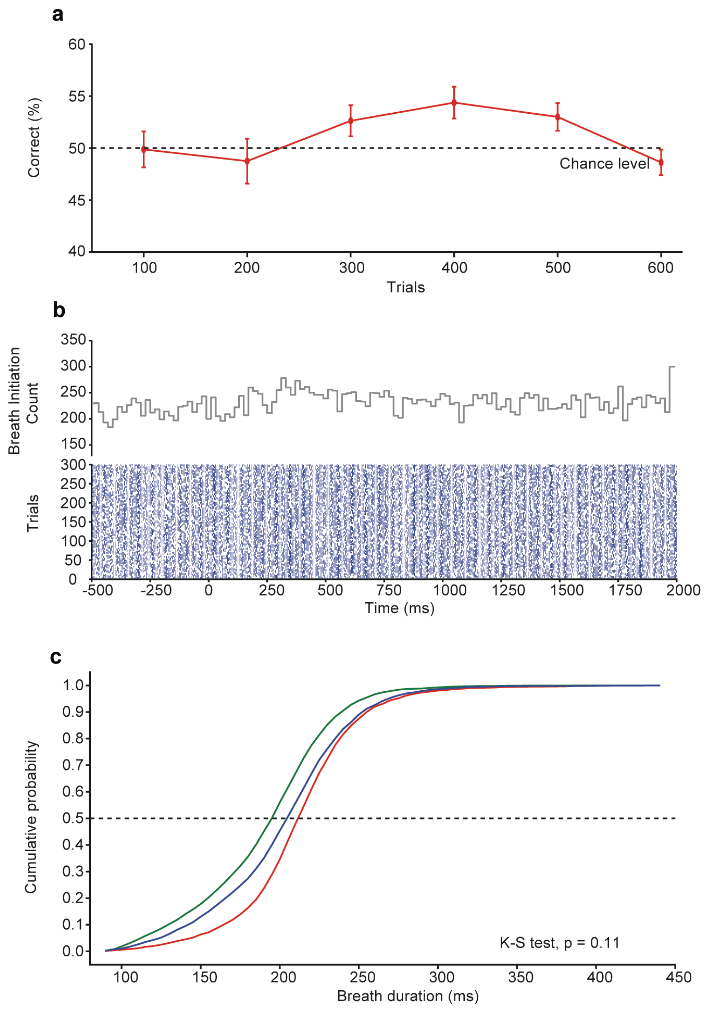


**Figure S3. Naïve mice show no specific sniffing peak latencies, Related to Figure 4**

(a) Animals showed no learning for a MO vs. MO discrimination task (ANOVA, F = 1.66, p = 0.16). Accuracy of discrimination is shown as % correct choices for 100 trials. Each data point is the average of 8 animals. The abscissa reflects progression of time. Data are presented as mean ± SEM.

(b) Raster plot and histogram, measured from the last 300 trials of MO vs. MO discrimination task (n = 8 mice).

(c) Cumulative probability distribution plot of breath duration measured for pre- (red), post- (blue) and during (green) decision-making period. Breath duration remains unaltered during the three phases (K-S test, p = 0.11).

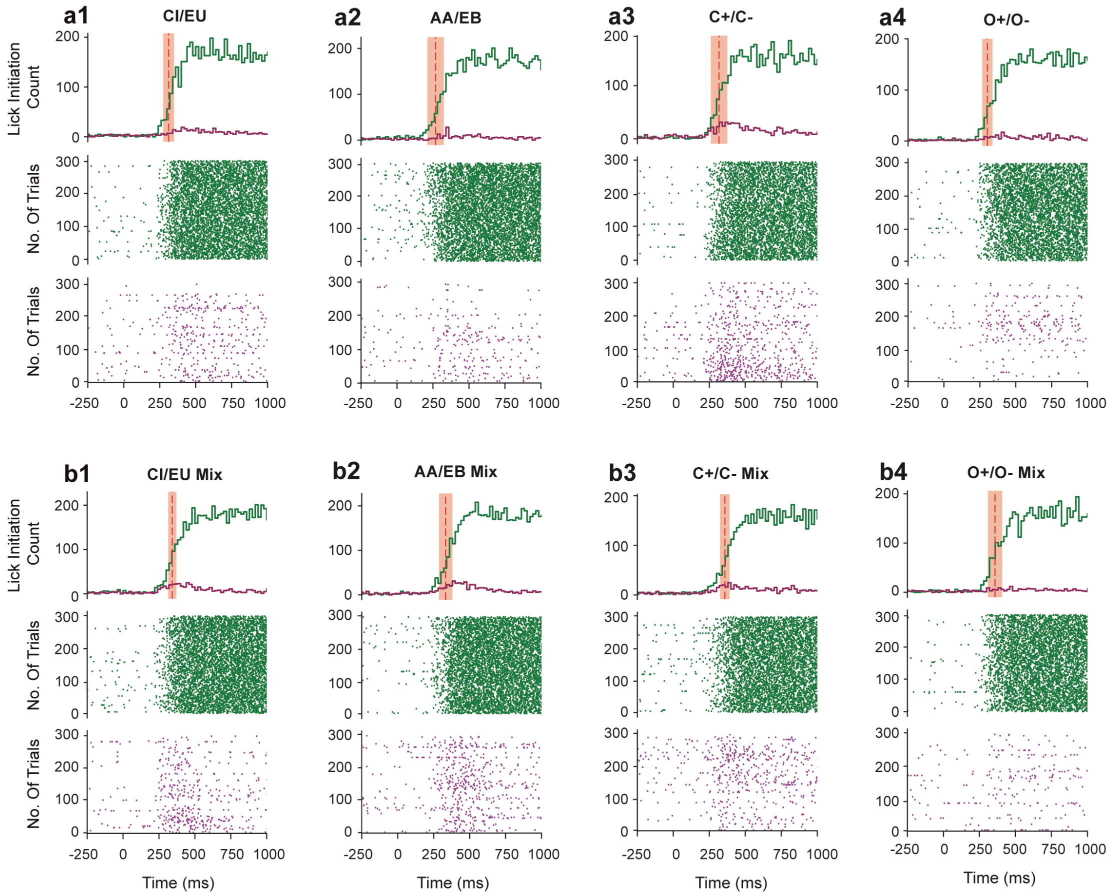


**Figure S4. Licking behavior of mice towards rewarded and non-rewarded odorants in a discrimination task, Related to Figure 4**

(a1-a4) Lick onset pattern for monomolecular odorants

(b1-b4) Lick onset pattern for binary mixtures

Raster plots (bottom two rows of each panel, a1 – b4, n = 7-8 mice) show the lick onset pattern of the mice during last task of training where the performance accuracy was ~90%. Each point on the raster represents the start of licking. Histograms were (top row of each panel, a1 – b4, n = 7-8 mice) calculated from all trials using a bin size of 20 ms (i.e. adding up the number of lick onsets initiated within 20 ms across all trials). The red dotted line and shaded region represent the mean ODT ± SEM. Green colored dots and histogram represents the rewarded trials and magenta dots and histogram represent non-rewarded trials. K-S test comparing Lick Initiation Count histograms of rewarded and non-rewarded trials; $p < 0.0001$ for all odor pairs. Lick responses during -250 to 1s are shown here, combined lick histogram of complete duration is shown in Figure 4.

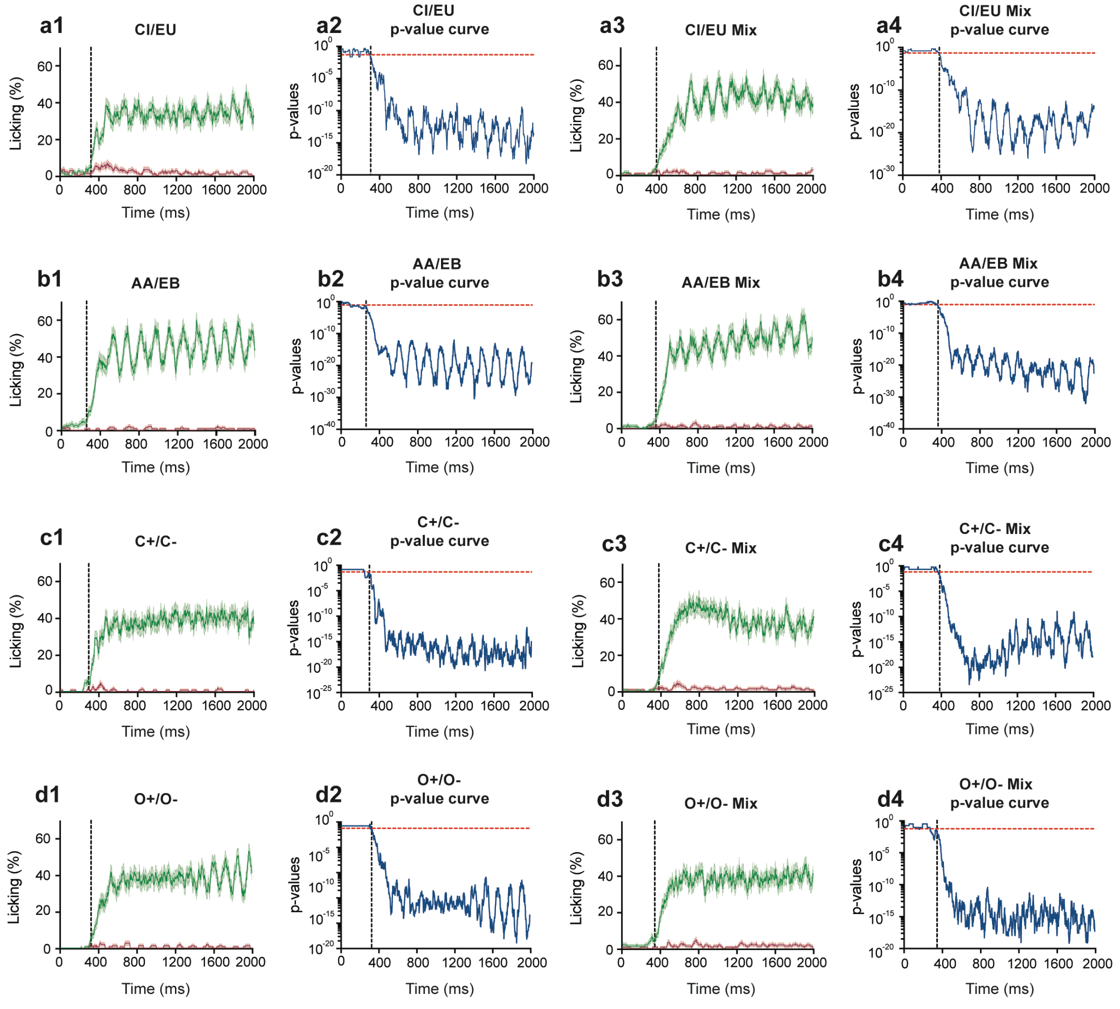


**Figure S5. Odor discrimination time measurements from the licking behavior of mice towards rewarded and non-rewarded odorants, Related to Figure 5**

(a1, b1, c1, d1) Averaged (150 trials, corresponds to one task) lick responses for simple odors observed for an individual representative animal. Green traces represent the licking responses for rewarded odorants and magenta traces represent non-rewarded trials.

(a2, b2, c2, d2) Statistical difference calculated between 150 trials of rewarded and non-rewarded trials. The red dotted lines indicate p-value of 0.05 and the black dotted line indicates where the traces cross the p-value of 0.05 and corresponds to the calculated ODT.

(a3, b3, c3, d3) Averaged lick responses for binary mixtures and (a4, b4, c4, d4) statistical difference calculated for binary mixtures.

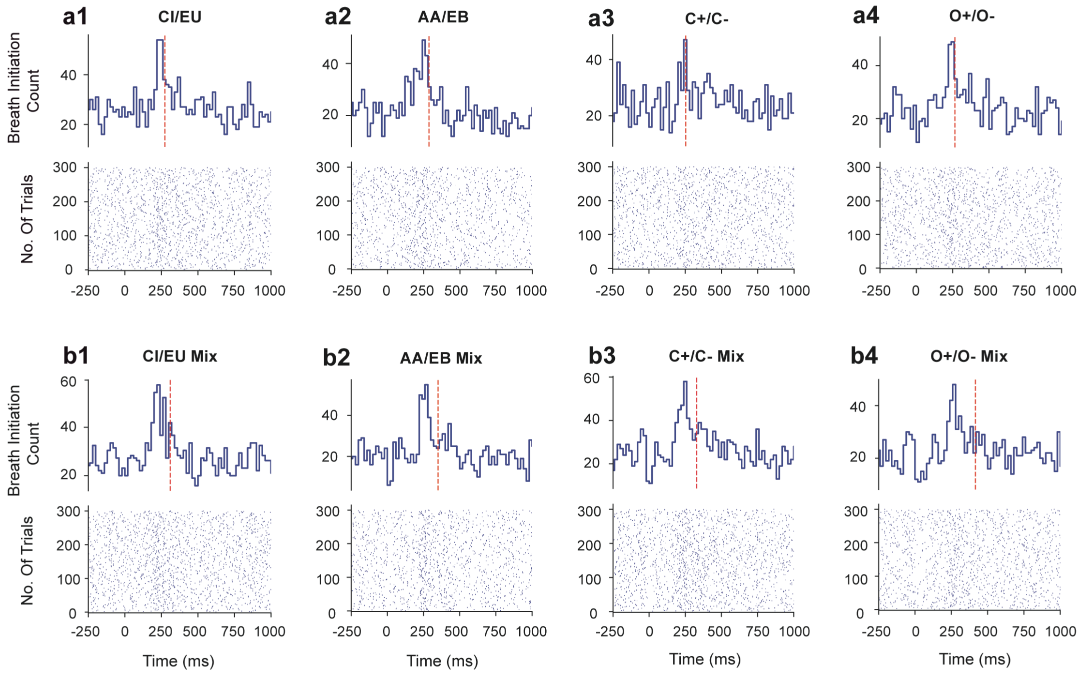


**Figure S6. SPL and ODT measurements from individual mice, Related to Figure 5**

(a1-a4) Breath initiation pattern for monomolecular odorants

(b1-b4) Breath initiation pattern for binary mixtures

Bottom half of each panel (raster plots of a1 – b4) shows the breathing cycle pattern of an individual mouse from a task of 300 where the performance accuracy was ~90%. Each point on the raster represents the start of inhalation. Top half of each panel (a1 – b4) shows a histogram taken from all trials using a bin size of 20 ms (i.e. adding up the number of breathing cycles initiated within 20 ms across all trials). The red dotted line represent ODT measured from the same task.

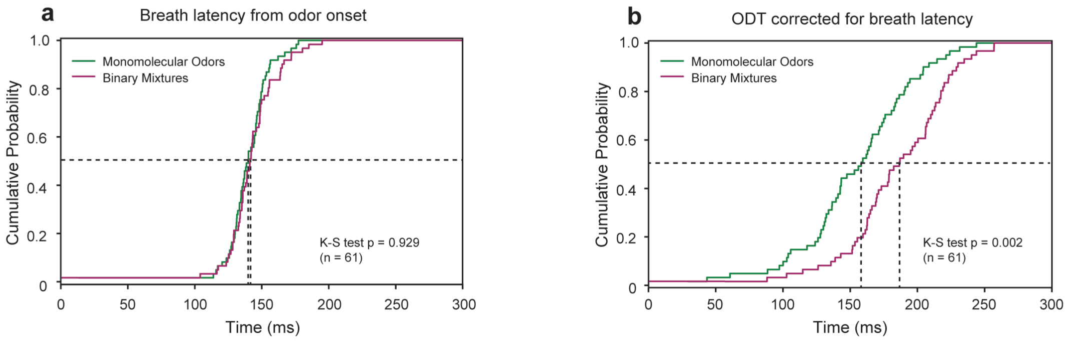


**Figure S7. Reaction time differences between simple and complex odors are independent of first breath onset delays, Related to Figure 5**

(a) Comparison between first breath onset delays between monomolecular odors and binary mixtures.

(b) Corrected ODT measurements after subtracting the first breath onset delay. Mice showed longer ODTs for binary mixtures compared to monomolecular odors.
